# Resolving heterogeneity of targeted lipid nanoparticles through solution-based biophysical analyses

**DOI:** 10.64898/2026.03.31.715590

**Authors:** Hannah C. Geisler, Hannah C. Safford, Ajay S. Thatte, Marshall S. Padilla, Elisa Battistini, Hannah M. Yamagata, Violet M. Ullman, Alex Chan, Benjamin E. Nachod, Anushka Agrawal, Maxwell B. Watkins, Jesse B. Hopkins, Andrew Tsourkas, Kushol Gupta, Michael J. Mitchell

## Abstract

Targeted lipid nanoparticles (tLNPs) represent the next frontier in nucleic acid therapeutics, enabling cell-specific delivery through covalent attachment of targeting ligands that drive receptor-mediated uptake. tLNPs are particularly promising for pregnancy-associated applications where precise on-target delivery is required to minimize maternal toxicity and protect fetal health. Yet, their rational design is limited by an incomplete understanding of how tLNP physicochemical properties influence biological performance. Conventional LNPs already exhibit pronounced heterogeneity in size, composition, and RNA loading, which is further amplified in tLNPs by variability in ligand attachment and surface density. Because traditional analytical methods report only ensemble-averaged properties, the nanoscale diversity of tLNPs remains unresolved. Here, we find that tLNP functional behavior is governed by previously inaccessible, structurally distinct tLNP subpopulations that are not captured by bulk measurements. We utilize asymmetric flow field-flow fractionation integrated with in-line UV spectral analysis, light scattering, and synchrotron small-angle X-ray scattering (AF4-UV-DLS-MALS-SAXS) to resolve ligand-dependent tLNP subpopulations that differ in size, shape, composition, and relative abundance. We find that protein conjugation preserves the internal lipid–RNA nanostructure of base LNPs but substantially increases particle heterogeneity, particularly for larger and multivalent targeting ligands. Despite increased heterogeneity, tLNPs functionalized with higher-avidity ligands achieve more effective targeted placental RNA delivery in mice, suggesting that binding avidity can offset the functional consequences of polydispersity. Chemometric SAXS analyses reveal that only SAXS-resolved tLNP subpopulations, not ensemble-averaged parameters, correlate with targeted placental transfection in vivo, whereas bulk-derived physicochemical metrics more strongly associate with nonspecific hepatic delivery. Together, this work harnesses a separation-coupled biophysical platform to resolve previously inaccessible tLNP subpopulations and demonstrates that subpopulation nanoscale structure, rather than bulk-averaged properties, dictates targeted RNA delivery. These insights provide a mechanistic foundation for rational engineering of next-generation precision targeted RNA LNP therapeutics.

## INTRODUCTION

Targeted lipid nanoparticles (tLNPs) are central to the advancement of precision nucleic acid therapeutics, enabling precise, cell-specific RNA delivery while minimizing systemic toxicity^1–3^. tLNPs are commonly generated through chemical ligation of protein targeting ligands to the LNP surface, facilitating receptor-mediated endocytosis of LNPs by specific cell types. Despite their promise, the rational design of tLNPs remains limited by an incomplete understanding of how tLNP physicochemical properties govern biological behavior. The base LNPs from which tLNPs are derived exhibit substantial heterogeneity in size, composition, and RNA loading^4–7^, and subsequent protein functionalization steps introduce additional variability in protein attachment and surface density, yielding complex product mixtures whose true nanoscale diversity remains unresolved^8^ (**Fig. 1a**). Consequently, the inability to directly resolve and quantify tLNP physicochemical complexity has emerged as a fundamental barrier to the design and deployment of targeted nucleic acid delivery platforms for in vivo applications that demand strict cell specificity, such as chimeric antigen receptor (CAR) T cell therapies^9–11^, gene editing strategies^12–15^, and gene therapies for obstetric complications^16–18^.

**Figure 1.**
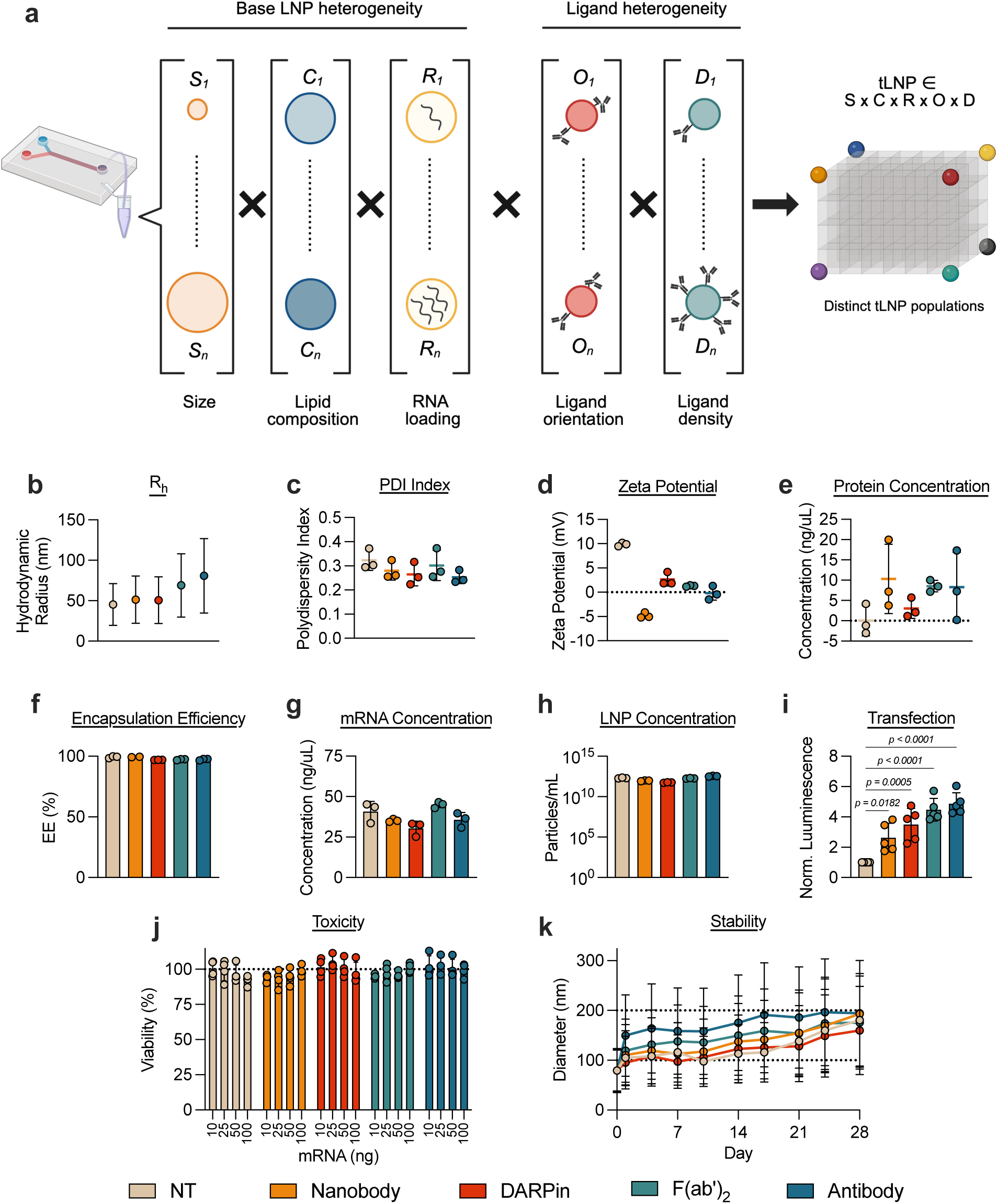
Heterogeneity of targeted LNPs (tLNPs) is unresolved by traditional bulk measurements. (**a**) Schematic depicting polydispersity intrinsic to base LNPs, where the addition of targeting ligands further amplifies this heterogeneity, resulting in a product space of infinite combinations. This image was created with BioRender.com. LNP physicochemical characteristics including (**b**) R_h_ and (**c**) PDI determined via DLS, (**d**) ζ-potential calculated via electrophoretic light scattering, (**e**) apparent protein concentration determined via a potassium cyanide (KCN) assay, (**f**) encapsulation efficiency and (**g**) mRNA concentration determined by a RiboGreen assay, and (**h**) LNP particle concentration determined via DLS. (**i–j**) NT LNPs and tLNPs containing FLuc mRNA were incubated with A431 cells at a dose of 25 ng of mRNA per 25,000 cells. (**i**) Luciferase mRNA delivery and (**j**) cell viability in A431 cells 24 h after treatment with nontargeted (NT) LNPs or anti-EGFR tLNPs. Normalized luciferase expression was quantified by normalizing to cells treated with NT LNPs. Percent cell viability for each treatment condition was normalized to untreated cells. (**k**) Stability of LNPs stored at 4°C in aqueous buffer for 28 days. Measurements are reported mean ± SD from *n* = 3 technical replicates for (**a–g), (k)** and *n* = 5 biological replicates for (**i–j).** One-way ANOVA with post hoc Student’s t-tests using the Holm–Sídak correction for multiple comparisons was used to compare luminescence in (**i)** across treatment group.

Common practices for (i) confirming successful protein functionalization and (ii) characterizing tLNP products rely upon bulk ensemble-average measurement, such as dynamic light scattering (DLS) and plate-based assays (BCA, fluorescent protein quantification). Traditionally, increases in LNP hydrodynamic radius (R_h_) measured by DLS has been interpreted as evidence of successful protein functionalization, while plate-based assays provide an estimate of the total protein incorporated into the final tLNP product. However, these approaches lack the resolution required to characterize heterogeneous tLNP populations^19,20^. For example, DLS cannot resolve subpopulations within polydisperse samples^21,22^, and calculated R_h_ measurements disproportionately reflect larger species^23^. Moreover, DLS assumes that particles are perfectly spherical, despite recent evidence that LNPs adopt ellipsoidal geometries in solution^24^. Similarly, plate-based protein quantification assays yield only bulk, ensemble-averaged measurements and cannot report on particle-level variability in ligand loading or spatial distribution^25,26^. Finally, none of these protein-detection methods report on internal LNP structure. Thus, existing analytical strategies obscure the true nanoscale structural and compositional landscape of tLNP formulations, limiting direct establishment of predictive structure–function relationships between protein functionalization patterns and biological outcomes.

Here, we leverage emerging solution-phase, “plug-and-play” orthogonal biophysical technologies to resolve the nanoscale architecture and compositional heterogeneity of tLNPs functionalized with a panel of protein targeting ligands differing in size and valency. Recent advances in separation-coupled biophysical platforms, including the integration of asymmetric flow field-flow fractionation (AF4) with synchrotron small-angle X-ray scattering (SAXS), have enabled nanoscale structural resolution of proteins^27,28^ and, more recently, mRNA LNPs^21,22,29–31^. However, these approaches have not been systematically applied to tLNPs, where protein functionalization introduces additional layers of compositional and structural heterogeneity, nor have they been extensively used to directly link tLNP physicochemical features to biological targeting capacity. In this work, we integrate AF4 with in-line UV–Vis detection, dynamic light scattering (DLS), multi-angle light scattering (MALS), and SAXS (AF4-UV-DLS-MALS-SAXS) to characterize tLNPs in solution. This approach enables separation-resolved analysis of co-existing particle populations and provides complementary measurements of composition (UV–Vis), hydrodynamic size (R_h_, from DLS), molar mass and radius of gyration (M_w_ and R_g_, from MALS), and nanoscale structure (SAXS). Through chemometric analysis of both UV–Vis spectra and SAXS scattering profiles, we demonstrate orthogonal resolution of previously inaccessible, distinct tLNP subpopulations and quantify their relative abundance, size, and structural features. Batch SAXS measurements establish that protein conjugation preserves the internal lipid–RNA organization of LNPs, while separation-coupled analyses uncover substantial surface-driven structural heterogeneity that is not accessible to ensemble methods. This integrated approach thus enables direct resolution of distinct tLNP subpopulations and their structural descriptors, including R_g_, maximum dimension (D_max_), and particle morphology.

We further demonstrate the functional relevance of this biophysical framework by evaluating relationships between tLNP physicochemical properties and RNA transfection in a murine model of pregnancy, a setting where maternal–fetal safety constraints are exceptionally stringent and effective therapeutic options for obstetric complications remain scarce^32,33^. By evaluating tLNP-mediated RNA delivery in vitro in placental cells and in vivo in pregnant and nonpregnant mice, we find that SAXS-resolved tLNP subpopulations show stronger correlation with targeted placental RNA delivery than ensemble-averaged measurements, whereas ensemble-averaged parameters instead predict nonspecific hepatic transfection and inflammatory responses. Together, this work demonstrates a separation-coupled, solution-based biophysical framework to resolve functionally relevant tLNP subpopulations, define their nanoscale structure, and link these features to biological outcomes, thereby enabling subpopulation-level structure–function relationships to guide the rational design of precision RNA tLNP therapeutics for pregnancy complications and other highly cell-specific in vivo applications.

## RESULTS

### Heterogeneity of tLNPs is unresolved by traditional bulk measurements

LNPs were formulated via chaotic mixing of an organic lipid phase and an aqueous mRNA phase using polydimethylsiloxane microfluidic devices^34,35^. The base LNP was formulated using the placenta-tropic Lipid A4^36^. LNPs were formulated encapsulating firefly luciferase (FLuc) messenger RNA (mRNA) and were subsequently functionalized with one of four targeting proteins (nanobody, DARPin, F(ab’)_2_ fragments, or antibody) via strain-promoted azide-alkyne cycloaddition (SPAAC) chemistry as previously described^37–39^ (**Supplementary Fig. 1).** All protein targeting moieties were directed against the epidermal growth factor receptor (EGFR), a receptor highly expressed in both murine and human placental tissue^38,40^.

First, traditional LNP characterization techniques, including dynamic light scattering (DLS), electrophoretic light scattering, and plate-based fluorescent assays were employed to evaluate LNP physicochemical properties. The base, nontargeted (NT) LNP had a hydrodynamic radius (R_h_) of ∼45 nm, a polydispersity index (PDI) of ∼0.3, and a neutral ζ-potential around ∼10 mV (**Fig. 1b-d, Supplementary Table 1**). As expected, protein functionalization resulted in an increase in R_h_ accompanied by changes in ζ-potential. Protein functionalization did not influence LNP PDI as measured by DLS. Functionalization with smaller protein ligands (nanobody and DARPin) increased R_h_ by ∼5-6 nm, whereas larger protein ligands (F(ab’)_2_ and antibody) produced greater increases in R_h_ (∼24–35 nm). DARPin and F(ab’)_2_ tLNPs exhibited reduced but still positive ζ-potential, while nanobody and antibody tLNPs displayed net negative ζ-potential values. Results from a potassium cyanide (KCN)-mediated protein quantification assay confirmed that no protein was present in solution in NT LNPs. All tLNPs had bulk protein concentrations ranging from ∼10–15 ng µL^-1^ (**Fig. 1e**). Changes in physicochemical properties, including R_h_ and ζ-potential, in combination with results from the KCN-mediated protein quantification assay, indicated that all tLNP groups were successfully functionalized with their respective protein. Protein functionalization did not impact LNP encapsulation efficiency or mRNA concentration. All LNPs had encapsulation efficiencies ranging from ∼95–99%, mRNA concentrations ranging from ∼30–45 ng µL^-1^, and LNP particle concentrations ranging from ∼10^11^–10^12^ particles per ml (**Fig. 1f-h**).

Finally, all tLNPs showed enhanced mRNA delivery relative to NT LNPs when screened in EGFR+ A431 control cells (**Fig. 1i**), further validating successful protein functionalization. Protein addition did not introduce any detectable cytotoxicity, as none of the LNP formulations exhibited toxicity across a range of mRNA doses (**Fig. 1j**). In stability studies, all LNPs maintained consistent size distributions for approximately ten days at 4°C in aqueous buffer before measurable changes were detected by DLS (**Fig. 1k**). Notably, NT LNPs displayed pronounced increases in size over time, consistent with aggregation, whereas tLNPs exhibited smaller shifts in hydrodynamic radius, suggesting improved colloidal stability.

### Evidence for heterogeneity in tLNPs by SEC-MALS and AUC

Because conventional characterization methods provide limited insight into LNP composition and structural heterogeneity, size-exclusion chromatography with multi-angle light scattering (SEC-MALS) was applied to confirm composition of NT LNPs and to probe changes in apparent size following protein functionalization. While DLS assumes sphericity, MALS, when coupled with UV and refractive index (RI) detection, provides greater quantitative information on LNP composition, mass, and spatial extent^41^. Moreover, pairing MALS with SEC enables elution-resolved determination of physicochemical properties, allowing interrogation of heterogenous systems that conventional SEC or DLS alone cannot access^42^.

UV chromatograms from SEC provide the relative abundance of species eluting over the course of the separation, with each sample displaying a predominant peak between 22–30 min corresponding to the bulk LNP population (**Supplementary Fig. 2a-g**). Molar mass (M_w_) profiles and nucleic acid fraction were determined leveraging 18-angle light scattering and RI detection, with UV measurements corrected for LNP Mie scattering as previously described^41^. Conjugate analysis enabled determination of mRNA concentration in NT LNPs, confirming the expected two-component lipid-RNA composition (∼58 MDa total, ∼2% RNA mass) (**Supplementary Fig. 2a-b**). However, because tLNPs incorporate a third material component (protein), the standard two-component conjugate model cannot be applied, preventing direct measurement of RNA content.

Assuming ∼90 sites of conjugation on a base NT LNP of ∼58 MDa, added protein mass of anywhere between 1.4 to 13.5 MDa could be expected, comprising anywhere from 2% to 19% of the total protein mass fraction (**Supplementary Table 2**). Mass profiles of NT LNPs displayed sloping molar masses, ranging from ∼50 to ∼100 MDa across the elution peak (**Supplementary Fig. 2a**), consistent with the inherent polydispersity of NT LNPs. Comparison of chromatographic profiles showed that tLNPs eluted earlier than NT LNPs (**Supplementary Fig. 2c**), indicating increased apparent LNP size after protein functionalization. Using mass-averaged *dn/dc* values assuming 100% protein-conjugation, tLNPs mass profiles were calculated and found to be comparable to NT LNPs (**Supplementary Fig. 2d-g**); discrete subpopulations were not evident. The observed broad, sloped mass profiles across the unimodal elution peaks of all LNPs suggests substantial heterogeneity that chromatography alone cannot fully resolve. This substantial heterogeneity was further substantiated by sedimentation/flotation velocity analytical ultracentrifugation (SV/FV-AUC) (**Supplemental Fig. 3a-c**), which was applied to determine whether these apparent subpopulations could be separated by their density properties. Although SV-AUC showed small increases in sedimentation coefficients for tLNPs relative to NT LNPs (**Supplementary Fig. 3d**), these shifts did not correspond to discrete, resolvable species, indicating that SV-AUC also lacked the resolution needed to fully deconvolve tLNP heterogeneity.

### Batch SAXS reveals conserved internal structure with increased tLNP structural variability

Traditional batch SAXS measurements were made to assess tLNP internal structure via their Bragg peak features, and to determine whether protein functionalization affects internal structure.

Although protein attachment is generally assumed not to influence internal lipid–RNA arrangement, this has not, to our knowledge, been directly demonstrated.

The SAXS scattering profiles of mRNA-loaded LNPs are distinguished by a characteristic first-order Bragg peak at scattering vectors where (q) ≈ 0.1–0.15 Å^−1^ (d ≈ 41.9–62.8 Å), reflecting a highly ordered internal lattice arising from RNA–lipid interactions^43,44^ (**Fig 2a, Supplementary Fig. 4a**-**b**). The Bragg region of each scattering profile was analyzed using multi-Lorentzian peak fitting^45^, which separates the sharp internal-ordering signal from disorder on larger length scales. This approach enabled quantitative extraction of peak width (full width at half maximum, FWHM), peak intensity (I_p_), and integrated area^45–47^ (**Supplementary Table 3**). NT LNPs exhibited a first-order Bragg peak at q ≈ 0.12 Å^−1^, whereas tLNPs displayed similar peaks at q ≈ 0.11 Å^−1^, consistent with the characteristic lattice spacing of lipid–RNA packing (**Fig. 2b**). The FWHM of the Lorentzian fit serves as a measure of structural coherence: narrower peaks correspond to well-defined ordering, whereas broader peaks correspond to more disordered correlations^48^. Across all formulations, the Bragg peaks remained narrow, indicating preservation of internal structure following protein functionalization. Specifically, NT LNPs exhibited a FWHM of ∼0.045 ± 0.002, while nanobody, DARPin, F(ab’)_2_, and antibody tLNPs showed FWHMs of ∼0.038 ± 0.004, ∼0.039 ± 0.001, ∼0.038 ± 0.004, and ∼0.038 ± 0.001, respectively. The comparison of the area of the two respective Lorentzian peaks in the fit provides a relative measure of the proportion of order to disorder in the internal structure. In all cases except the antibody tLNP sample, these proportions are similar. The proportion of disorder is more pronounced in the antibody tLNP formulation, however, it is not clear how much of this is due to elevated heterogeneity.

**Figure 2.**
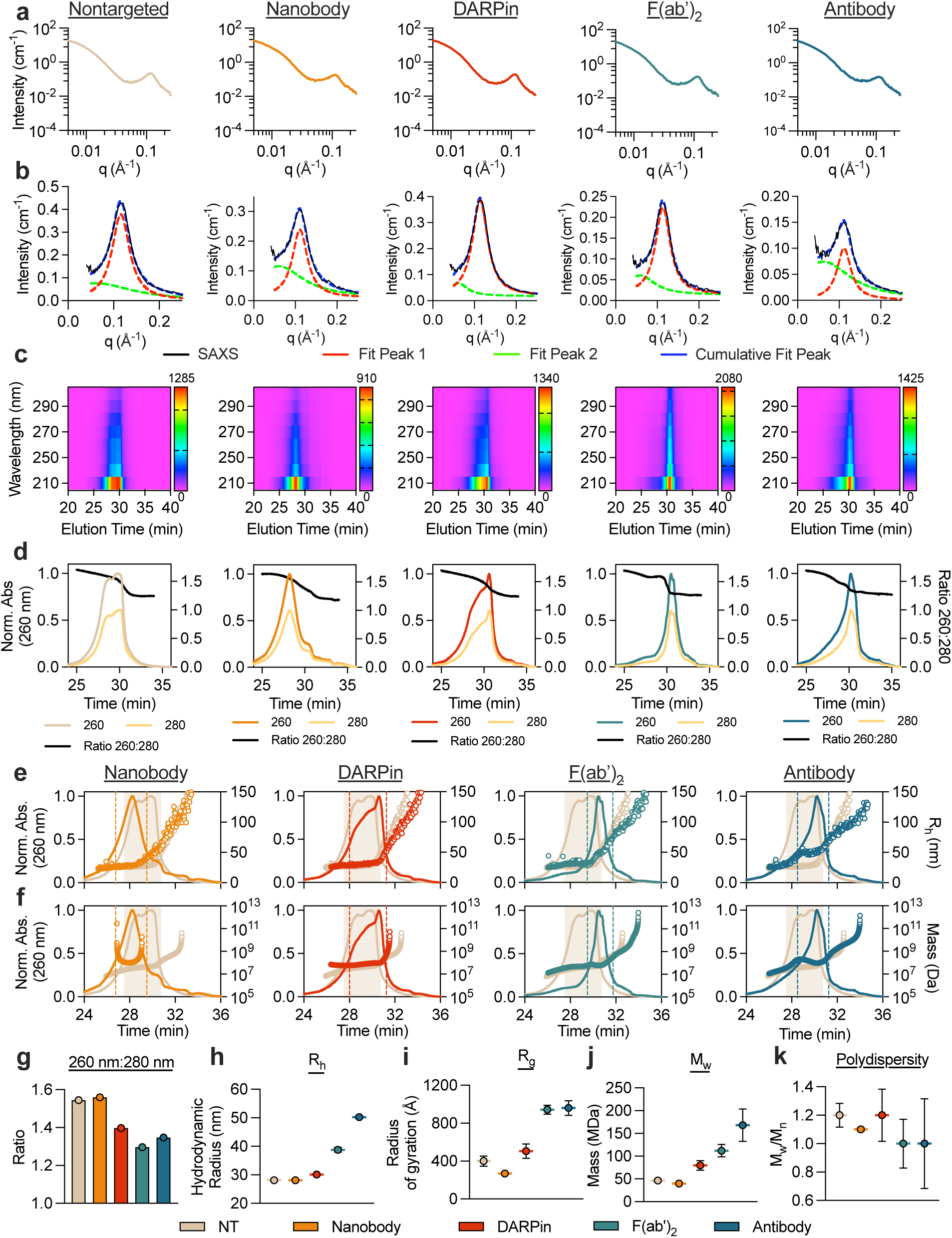
AF4-UV-DLS-MALS-SAXS resolves tLNP ligand-dependent polydispersity and compositional heterogeneity. (**a**) SAXS profiles of NT LNPs and tLNPs, showing the characteristic Bragg peak feature associated with internal lipid–RNA organization. (**b**) Fitting of the Bragg peak feature using a multiple-Lorentz model where red is the first-order Bragg peak fit, green represents higher-order particle disorder and blue is the cumulative fit of the two features. (c) In-line AF4-UV-Vis contour maps (200–300 nm) depicting wavelength-resolved absorbance of LNPs during separation. The 260 nm absorbance signal is characteristic of encapsulated RNA, enabling identification of RNA-containing LNP populations across the AF4 elution profile. Overlapping spectral features indicate the presence of multiple co-eluting populations with distinct compositional profiles, motivating subsequent chemometric deconvolution. (**d**) Absorbance at 260 nm (RNA-associated signal) and 280 nm (protein-associated signal) from (**c**) plotted against the corresponding 260:280 ratio for each LNP formulation. Deviations in the 260:280 ratio across elution time indicate heterogeneity in RNA and protein content, suggesting the presence of compositionally distinct subpopulations that cannot be resolved by bulk measurements alone. These data were further subjected to chemometric analysis (see **Supplemental Figure 6**). The determined (**e**) R_h_ profiles derived from in-line DLS and (**f**) molar mass profiles derived from MALS analysis for NT LNPs (beige) and tLNPs (colors) overlaid with UV fractograms from in-line AF4 separation. (**g**) Peak 260:280 ratios from (**d**), shown for comparison across LNP groups. (**h**) In-line DLS R_h_ and MALS-derived (**i**) mass, (**j**) radius of gyration, and (**k**) polydispersity plotted for comparison across LNP groups. Measurements are reported mean ± standard error for (**h–k).**

While Bragg peak features remain well conserved throughout the samples, the lower-q Porod region of the scattering profiles showed marked differences when superposed^47^ (**Supplementary Fig. 4c**). Although the Porod regime is sensitive to interfacial characteristics and surface heterogeneity, in ensemble-averaged batch SAXS these features reflect convolution of heterogeneous particle populations and therefore cannot be unambiguously assigned to specific structural species. As such, robust interpretation of these parameters is limited in batch measurements by sample heterogeneity.

### AF4-UV-DLS-MALS-SAXS resolves ligand-dependent tLNP polydispersity and compositional heterogeneity

To better resolve and study tLNP subpopulations, an analytical platform comprised of AF4 separation in-line with multiple modalities of biophysical analysis was applied: (i) in-line UV-Vis diode array detection, (ii) DLS, (iii) MALS, and (iv) synchrotron SAXS (**Supplementary Fig. 5).** This AF4-UV-DLS-MALS-SAXS approach leverages the high-flux, low-q capabilities of an advanced synchrotron source with the superior size resolution of AF4, enabling hydrodynamic separation and structural analysis to be performed in-line^21^. While SEC-SAXS has been used to characterize LNP systems^24^, AF4-SAXS has more recently emerged as a powerful approach for resolving the structural features of standard mRNA LNP formulations^21,29–31,49^. Because SEC separation is constrained by pore accessibility, LNPs and higher-mass assemblies are expected to elute near the void volume with limited resolution. In contrast, AF4 separates particles based on diffusion under an applied crossflow, enabling size-resolved analysis of nanoparticle-scale species without the exclusion limitations inherent to SEC. Building on these recent advances, the present work represents, to our knowledge, the first application of integrated AF4-UV-DLS-MALS-SAXS for real-time, in-line characterization of complex tLNP formulations. AF4 provides gentle, size-based fractionation, while UV-Vis, DLS, MALS, and SAXS collectively report compositional, hydrodynamic, and structural parameters across eluting fractions. Integrating all four modalities of analysis in-line enables a comprehensive view with orthogonal validation of tLNP heterogeneity in solution otherwise inaccessible to individual measures and bulk measurements.

AF4–UV fractograms provided substantially greater resolving power than SEC, revealing complex elution behavior across all LNP formulations. All samples exhibited a dominant peak between ∼28–32 min corresponding to the primary LNP population, accompanied by wavelength-dependent variations indicative of compositional heterogeneity. UV–Vis heat maps showed strong absorbance at 210 nm across all samples, consistent with lipid-dominated absorbance and scattering, along with a narrower feature at 260 nm associated with encapsulated RNA **(Fig. 2c)**. Absorbance at 280 nm increased progressively from NT LNPs to antibody-functionalized tLNPs, reflecting increasing protein incorporation. Analysis of the 260 nm trace revealed a bimodal RNA-associated peak for NT LNPs, suggesting the presence of at least two RNA-enriched subpopulations **(Fig. 2d–f)**, whereas tLNPs exhibited broader, asymmetric, and largely unimodal profiles consistent with overlapping species. Correspondingly, the 260:280 absorbance ratio decreased systematically from NT to antibody tLNPs, consistent with increasing protein content **(Fig. 2g)**. Together, these wavelength-dependent changes indicate that the AF4 elution profiles contain overlapping LNP populations that differ not only in size, but also in RNA-to-protein composition, motivating subsequent chemometric decomposition of both the UV–Vis and SAXS datasets.

To more rigorously resolve these overlapping contributions, multivariate chemometric analysis was applied to the full AF4–UV diode array dataset (**Supplementary Fig. 6**). Singular value decomposition (SVD)^50^ and evolving factor analysis (EFA)^51^ indicated the presence of compositionally distinct components within each fractogram, exceeding what could be inferred from individual wavelength traces alone. Multivariate curve resolution (MCR-ALS) decomposed the data into resolved concentration profiles and corresponding UV–Vis spectra, enabling separation of lipid-dominated, RNA-associated, and protein-associated components across the elution time domain. These results confirm that NT LNPs contain at least two RNA-enriched populations with distinct elution behavior, while protein-functionalized tLNPs display broader, partially overlapping populations characterized by increased protein contribution and reduced RNA-to-protein ratios. Although RNA and protein together comprise a minor fraction of total LNP mass, their strong and spectrally distinct UV absorbance enables robust resolution by multivariate analysis. Absolute absorbance intensities, particularly at shorter wavelengths, may be influenced by lipid-derived UV Mie scattering and should therefore be interpreted in terms of relative optical contributions rather than direct mass fraction. Importantly, these UV-derived components define the compositional heterogeneity present across the AF4 elution window and provide the basis for interpreting the SAXS-resolved components as occurrences of overlapping LNP populations that differ in RNA and protein content.

Elution behavior and variations in R_h_ by in-line DLS and R_g_ and mass by MALS were consistent with the compositional heterogeneity identified by UV chemometric analysis, indicating that the overlapping RNA- and protein-associated populations also differ in hydrodynamic size and mass. In contrast to SEC, where larger species co-elute near the exclusion limit, AF4 separates particles based on diffusion, such that larger species elute later. Accordingly, three of four tLNP formulations eluted later than NT LNPs, consistent with increased particle sizes (**Fig. 2e**). Nanobody and DARPin tLNPs showed modest R_h_ increases but clear increases in R_g_ and molar mass (M_w_), consistent with their relatively small (<20 kDa) protein ligands (**Fig. 2f,h-j, Supplementary Fig. 7**). In contrast, F(ab’)_2_ and antibody tLNPs exhibited continuous increases in R_h_ across the elution peak, reflecting substantial heterogeneity and access to size regimes that would be partially excluded in SEC-based separations. Antibody tLNPs displayed an R_h_ plateau followed by a secondary increase, suggesting a more uniform subpopulation followed by a larger, potentially aggregated population. F(ab’)_2_ tLNPs exhibited the broadest size distribution, with R_h_ values spanning ∼30–70 nm. All tLNPs contained minor populations of large particles, providing a tractable explanation for why bulk DLS measurements yield overestimated hydrodynamic radii relative to the R_h_ distributions observed across AF4 fractograms. Consistent with these size distributions, MALS-derived R_g_ and molar mass increased substantially across all tLNPs (**Fig. 2i-j**). DARPin and F(ab’)_2_ tLNPs exhibited mass increases of ∼50 MDa, while nanobody and antibody tLNPs increased by ∼100 MDa (**Fig. 2j**). Steep, sloping R_h_ and mass trajectories, combined with broad asymmetric elution peaks, suggest that F(ab’)_2_ and antibody tLNPs exhibit the greatest polydispersity, corroborated by M_w_/M_n_ values (**Fig. 2k**). In-line DLS and AF4-MALS derived physicochemical parameters are summarized in **Supplementary Table 4**.

Taken together, the AF4–UV data indicate that multiple LNP populations with distinct RNA-to-protein composition co-elute within the main fractogram peak. This compositional heterogeneity provides a direct rationale for subsequent SAXS chemometric decomposition, as the ensemble scattering signal arises from these overlapping populations. In this framework, UV chemometrics resolves compositional contributions across the elution window, while SAXS chemometrics resolves the corresponding structural contributions. Although baseline resolution is not achieved, AF4-MALS provides substantially improved separation of heterogeneous mixtures compared to SEC-MALS and DLS alone. Together, these data indicate that AF4-resolved tLNP heterogeneity is manifested across composition, size, and mass, enabling linkage of compositional variability to structural heterogeneity that is obscured in ensemble-averaged measurements.

### SAXS component analysis reveals distinct structural tLNP subpopulations

As indicated by UV and light scattering analyses (**Fig. 2**), the AF4 elution profiles showed broad multimodal peaks for all formulations (**Fig. 3a**), reflecting overlapping LNP populations that differ in RNA and protein content and that cannot be resolved by elution time alone. Differences in peak width and shape were consistent with the hydrodynamic heterogeneity observed with in-line light scattering, with F(ab’)_2_ and antibody tLNPs exhibiting the broadest and most asymmetric elution profiles. To resolve structural subpopulations contributing to the SAXS signal across the AF4 elution window, we performed component analysis using SVD and EFA as implemented in REGALS^52,53^. SVD was first used to estimate the number of statistically significant scattering components present; this analysis indicated that the SAXS signal cannot be adequately described by a single particle population. EFA then identified the elution boundaries over which each component contributed measurably to the total scattering, thereby identifying partially overlapping populations contributing to the scattering signal. Finally, component profiles were extracted to obtain physically consistent scattering curves for each subpopulation^53^. This approach enabled robust separation of overlapping signals and yielded physically consistent SAXS profiles corresponding to co-existing LNP subpopulations rather than purely mathematical decompositions. In this framework, UV chemometrics and light scattering analysis define compositional heterogeneity across the elution window, while SAXS chemometrics resolves the corresponding structural heterogeneity arising from the same AF4-resolved populations.

**Figure 3.**
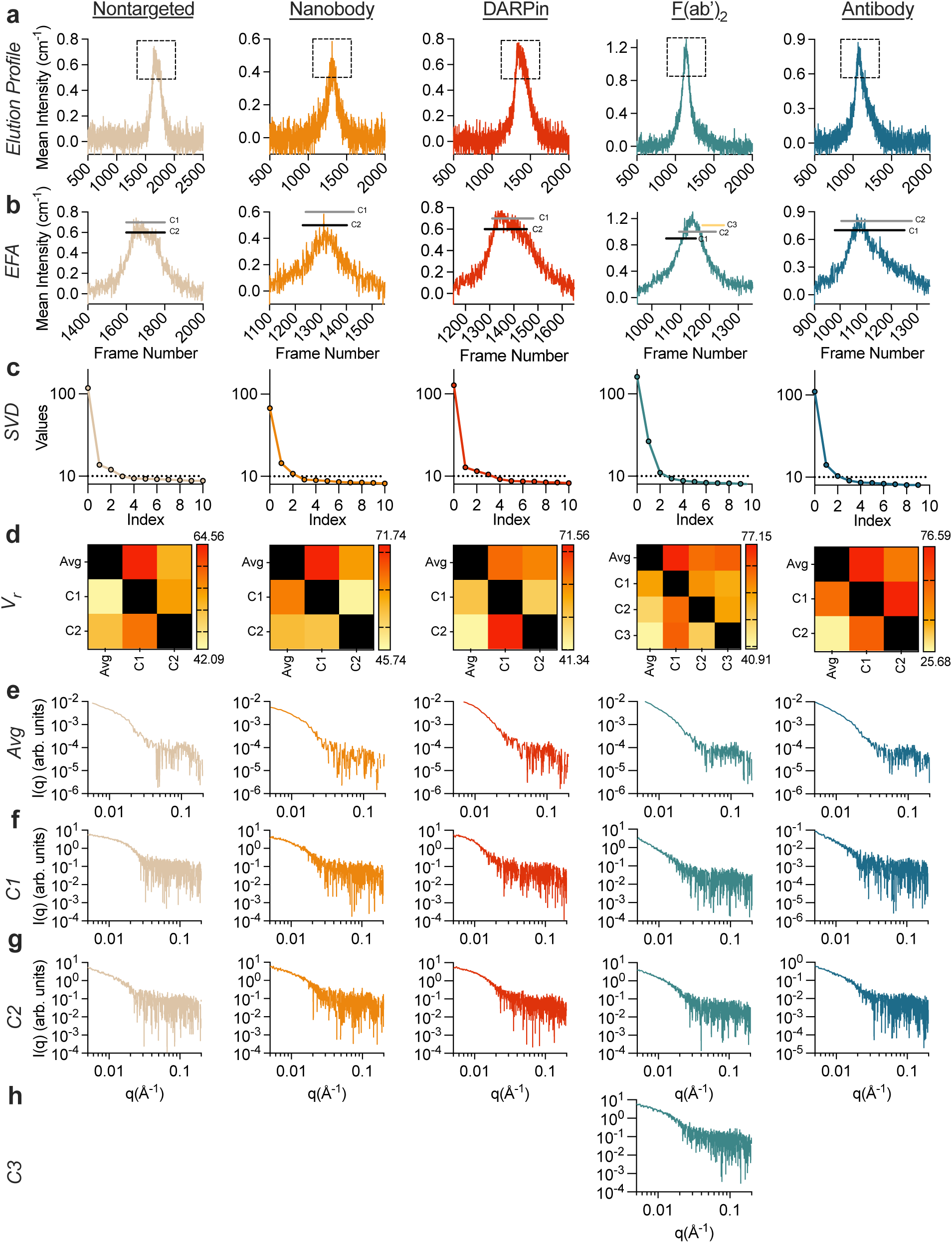
SAXS component analysis reveals discrete tLNP subspecies. (**a**) SAXS elution profiles of NT LNPs and tLNPs. Dotted boxes indicate the main LNP elution region, expanded in (**b**). Evolving factor analysis (EFA) (**b**) applied to AF4-SAXS data, highlighting frames corresponding to discrete evolving species (C1–C3) within each formulation. The resolved components correspond to distinct LNP subpopulations that differ in size and internal organization and a third component (when present) reflecting a minor population of highly heterogeneous or higher-mass particles. The temporal evolution of EFA-selected frames across the elution peak indicates that these components arise from partially overlapping populations that co-elute but differ in hydrodynamic size and composition. (**c**) Singular value decomposition (SVD) of SAXS datasets, indicating the number of independent signals present in each formulation. The presence of multiple significant singular values indicates that the SAXS signal cannot be described by a single particle population, necessitating decomposition into independent scattering components. (d) Volatility of ratio (V_r_) analysis used to assess the statistical uniqueness of deconvoluted component subpopulations relative to the ensemble-averaged population. (**e-h**) Regularized alternating least squares (REGALS) deconvolutions of SAXS data yields component-resolved scattering profiles. Shown are the (**e**) ensemble-averaged SAXS profile over the elution peak, and the REGALS-derived scattering profiles for the (**f**) C1, (**g**) C2, and (**h**) C3 subpopulations. These structural components are consistent with compositionally distinct populations inferred from UV-based chemometric analysis, linking RNA-associated heterogeneity to differences in particle size and morphology. Together, these analyses demonstrate that the ensemble SAXS signal arises from multiple, structurally distinct tLNP subpopulations that can be resolved and interpreted through chemometric decomposition.

EFA identified evolving scattering contributions across each elution peak (**Fig. 3b**). In the selected ranges, all formulations exhibited at least two evolving populations, except for F(ab’)_2_ LNPs, which exhibited three evolving subpopulations, which is consistent with the increased compositional and structural heterogeneity inferred from UV and light scattering measurements. The EFA-defined regions were broader for DARPin, F(ab’)_2_, and antibody tLNPs, consistent with greater size and mass heterogeneity. SVD quantified the number of statistically significant scattering components present in each dataset (**Fig. 3c**). In all samples, the first component (C1) dominated the signal, indicating one principal structural population. All LNP formulations showed non-negligible secondary components (C2), and for F(ab’)_2_ tLNPs, a non-negligible third component (C3), suggesting the presence of minor but distinct structural contributors in all formulations.

The volatility of ratio (V_r_) metric was then used to substantiate the statistical uniqueness of the SAXS data (**Fig. 3d, Supplementary Fig. 8**). The V_r_ parameter reports the variability in the ratio of scattering intensity between two profiles, quantifies differences between scattering profiles across the q-range selected, and is advantageous relative to other methods of profile comparison, especially in regimes of weaker signals^54^. In V_r_ matrices, lower V_r_ values (yellow) represent more similar profiles and higher V_r_ values (red) represent more distinct curves. V_r_ matrices of tLNPs revealed structurally distinct scattering profiles between the first, second, and third components (C1-C3), supporting the interpretation that these components represent distinct scattering subpopulations rather than numerical artifacts. These results support that the resolved components are statistically independent and non-redundant, rather than arising from overfitting of the SAXS data. Higher V_r_ values observed for F(ab’)_2_ and antibody tLNPs reflect comparatively increased differences when compared to the components of the other particles.

V_r_ matrices provided additional insight into structural differences among the resolved SAXS components. For NT LNPs, C1 most closely resembled the peak-averaged scattering profile, indicating that it is most consistent with the dominant structural contribution. In contrast, for nanobody, DARPin, and antibody tLNPs, the C2 showed the closest correspondence to the average profile, suggesting a shift in the dominant structural contribution following protein functionalization, indicating that the relative contribution of LNP subpopulations shifts with ligand incorporation. F(ab’)_2_ tLNPs exhibited the greatest heterogeneity: C3 best matched the peak-averaged profile, followed by the C2, while C1 was the most structurally distinct.

Across all extracted component scattering profiles (**Fig. 3e–h**), Bragg features were not sharply defined. This is likely due to peak broadening and sample dilution inherent to the in-line AF4-SAXS workflow and does not indicate a loss of internal lipid–RNA ordering, which was supported by batch SAXS measurements. Instead, the REGALS-derived profiles emphasize differences in the low-q and mid-q regions, reflecting variations in particle size and polydispersity among components, where low-q differences primarily reflect size and mass distribution, and mid-q differences reflect internal structural organization. These differences were most pronounced for F(ab’)_2_ and antibody tLNPs, reinforcing the conclusion that larger targeting ligands produce broader distributions of nanoparticle morphologies.

The high-q region of each SAXS profile was then analyzed using Porod scaling to probe short-length-scale density correlations. In contrast to ensemble-averaged batch SAXS, where Porod behavior reflects convolution of heterogeneous populations, AF4- and SVD-resolved profiles enabled interpretation of Porod scaling at the subpopulation level (**Supplementary Table 5**). Porod exponents for all LNP formulations fall predominantly in the range where m ≈ 3.0–3.8, consistent with compact particles exhibiting surface-fractal scattering rather than ideal Porod behavior. This is consistent with rough or diffuse interfaces, which would be expected due to lipid heterogeneity, heterogenous RNA–lipid organization within LNP interiors, and heterogenous protein functionalization. No formulation displays sustained mass-fractal behavior, although isolated SVD-resolved components approach m ≈ 2.5, consistent with locally disordered internal domains rather than globally fractal particles.

### Determination of model-independent structural parameters of distinct tLNP subpopulations

Having resolved the principal scattering components for each formulation, we next quantified the underlying structural properties for these subpopulations. Classical Guinier analysis, which directly ascertains the particle radius of gyration (R_g_) from the lowest q regime of recorded scattering, is often challenging for LNPs because their large real-space dimensions typically require low measurable q that is inaccessible in standard experimental configurations^24,55,56^. As a result, most LNP SAXS datasets lack a true Guinier region and may not always provide reliable R_g_ values depending on q-range and sample heterogeneity. However, the low-q coverage afforded by this AF4-SAXS experimental configuration enabled Guinier analysis for components that satisfied standard linearity criteria. Importantly, these Guinier analyses were performed on component-resolved SAXS profiles, reducing the impact of population heterogeneity that typically obscures low-q behavior in ensemble measurements. By combining AF4 separation with SAXS component analysis, these measurements enable structural parameters to be assigned to subpopulations that would otherwise be obscured in ensemble-averaged scattering data.

For NT, nanobody, and DARPin LNPs, both C1 and C2 components produced linear Guinier regions (**Fig. 4a**). For NT LNPs, C1 yielded an R_g_ of 132.4 ± 18.4 Å, while C2 exhibited a larger R_g_ of 284.6 ± 8.6 Å (**Supplementary Table 6**). Nanobody tLNP C1 and C2 had R_g_ values of 188.6 ± 8.0 Å and 219.9 ± 13.4 Å, respectively, and DARPin tLNPs produced comparable R_g_ values of 175.9 ± 11.6 Å and 177.3 ± 7.7 Å. For the F(ab’)_2_ and antibody tLNPs, only C3 and C2, respectively, had measurable Guinier regions; additional components could not be analyzed by Guinier methods due to their large size or the absence of a well-defined low-q linear regime within the measured range. The F(ab’)_2_ tLNP C3 component yielded an R_g_ of 190.4 ± 18.1 Å, while the antibody tLNP C2 exhibited an R_g_ of 232.9 ± 18.4 Å. The observed increases in R_g_ are consistent with both larger particle dimensions and broader mass distributions in protein-functionalized LNPs. These differences in R_g_ across subpopulations are consistent with the presence of structurally distinct subpopulations identified by SAXS chemometric analysis and with compositional heterogeneity inferred from UV and light scattering measurements.

**Figure 4.**
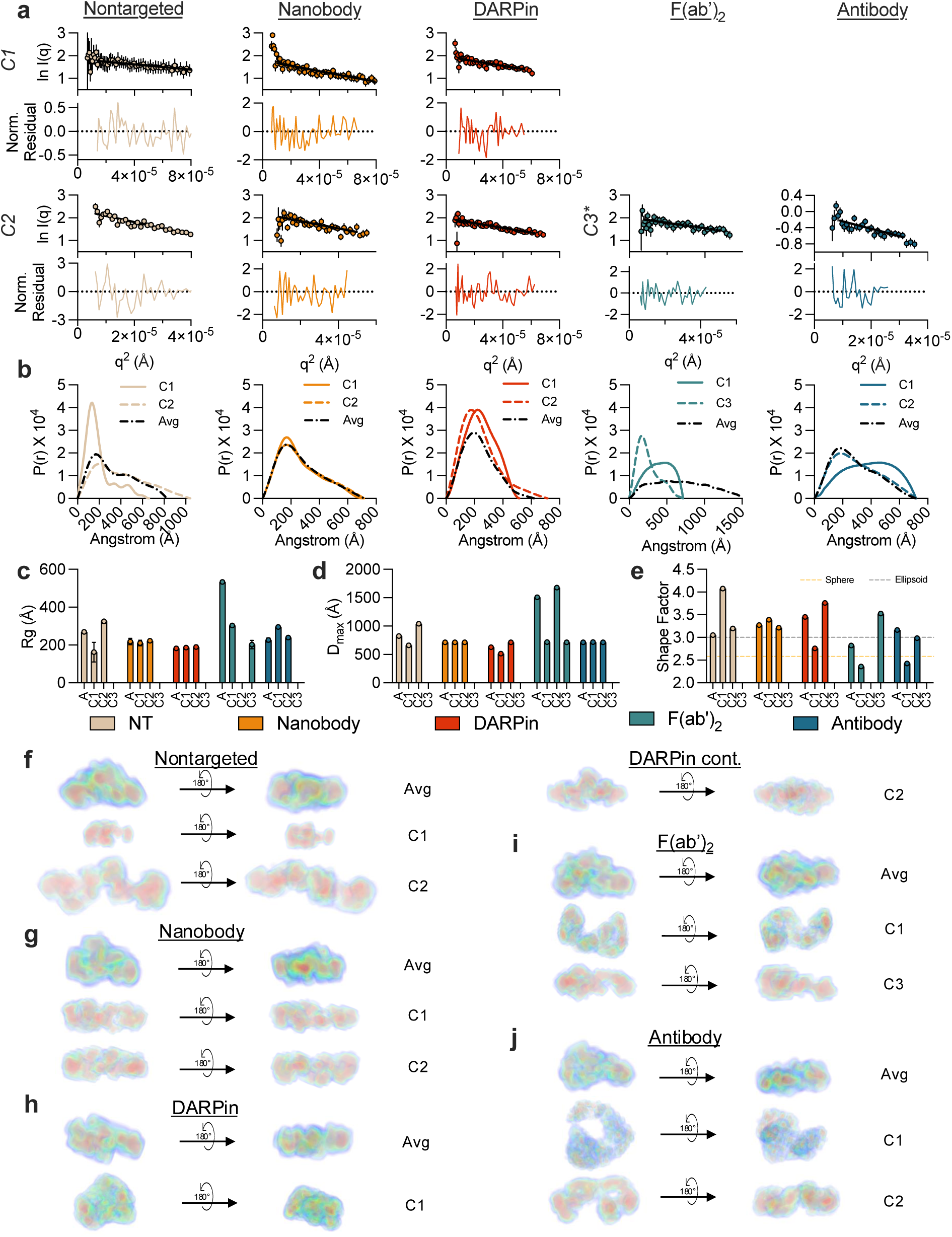
Determination of model-independent structural parameters of distinct tLNP subspecies. (**a**) Guinier analyses with corresponding residuals for SVD-resolved C1 and C2 components of NT LNPs and tLNPs, with the exception of F(ab’)_2_ tLNPs, where only C3 is shown. White regions indicate components for which Guinier analysis failed due to large size (q_min_R_g_ > 1.3). (**b**) P(r) analyses normalized by I(0) for the average profile and individual components (C1–C3) for NT LNPs and tLNPs. (**c**) Radius of gyration (R_g_) and (**d**) maximum dimension (D_max_) of the average profile and individual components derived from GNOM analysis for NT LNPs and tLNPs. Blank regions denote populations where GNOM analysis was invalid (q_min_D_max_ > 4). (**e**) LNP shape factor calculated as D_max_/ R_g_, where values of ∼2.58 and ∼3.0 correspond to spherical and prolate ellipsoid geometries, respectively. DENSS ab initio electron density reconstructions from the AF4-UV-DLS-MALS-SAXS profiles for (**f**) NT LNPs, (**g**) nanobody tLNPs, (**h**) DAPRin tLNPs, (**i**), F(ab’)_2_ tLNPs, and (**j**) antibody tLNPs.

The P(r) function represents the distribution of all intraparticle distances and is used to extract the real-space size parameters R_g_ and the maximum dimension (D_max_), providing complementary, global size estimates that are less sensitive to limitations in Guinier analysis and provide a more complete description of particle size distributions^56^. NT, nanobody, and DARPin LNPs exhibited more compact, unimodal P(r) curves, consistent with smaller ellipsoids (**Fig. 4b-d**). F(ab’)_2_ and antibody tLNPs displayed broader P(r) profiles with longer maximum dimensions, consistent with greater structural heterogeneity. All parameters derived from Guinier analysis and P(r) analysis using GNOM are summarized in **Supplementary Table 6**.

We also calculated a shape factor (D_max_/R_g_) for each component (**Fig. 4e, Supplementary Table 7**), where ideal spheres have values of 2.58 and prolate ellipsoids have values of 3.0^24,57^. NT LNPs exhibited shape factors consistent with ellipsoidal geometries, in agreement with previous reports of LNP morphology^24,58^. In contrast, tLNPs trended downward in shape factor with increasing ligand size, approaching more isotropic values. This shift may reflect preferential protein decoration on the elongated axes of ellipsoidal particles, effectively “rounding” their apparent shape by evening out surface mass distribution. Finally, the algorithm density from solution scattering (DENSS) can be applied to LNP SAXS data to generate low-resolution electron density reconstructions consistent with the scattering data^58^ (**Fig. 4f-j**). Consistent with trends observed in P(r) analyses and previous reports of shape, NT LNPs were anisotropic ellipsoids. To define length, width, and height independent of the arbitrary initial orientation of DENSS reconstructions^59^, density maps were aligned to their principal axes prior to dimensional analysis. Determined dimensions from DENSS showed that tLNPs varied substantially in their reconstructed dimensions across subpopulations (**Supplementary Table 8**). Relative to NT LNPs, all targeted formulations exhibited measurable shifts in one or more principal axes, consistent with ligand-dependent differences in particle morphology. Larger ligands (F(ab’)_2_ and antibody) were associated with components exhibiting greater size variability, with some becoming more elongated while others approached more isotropic, sphere-like dimensions, consistent with increased structural heterogeneity inferred from SAXS component analysis and the previous changes observed in the shape factor.

### Biophysical-biological correlations identify structural subpopulations governing tLNP targeting to the placenta

Designing effective tLNP nucleic acid delivery systems requires parameters that reliably predict delivery outcomes, yet such predictors remain largely undefined due to limited analytical resolution. To determine whether high-resolution biophysical measurements can fill this gap, we evaluated our tLNP library in vitro in immortalized placental cells and in vivo in pregnant mice and nonpregnant mice. All targeting ligands used in this study were anti-human EGFR proteins. Because human and mouse EGFR differ in their binding interfaces, the anti-EGFR protein ligands used in this work display negligible affinity for endogenous mouse EGFR, except in tissues with extremely high receptor density^60^. The placenta, which expresses abundant EGFR and is directly exposed to maternal circulation^38,61,62^, therefore represents the primary tissue in which ligand-receptor engagement would be expected. This design ensured a model where pregnancy-specific targeting could be cleanly resolved from baseline biodistribution.

When screened in human placental trophoblasts, antibody tLNPs showed the greatest early (1 h, 4 h timepoints) cellular accumulation compared to NT LNPs and all other tLNP groups (**Fig. 5a-b, Supplementary Fig. 9**). By 24 h, antibody tLNP accumulation remained significantly more enriched than NT LNPs, however both F(ab’)_2_ and antibody tLNPs produced significantly higher mCherry expression relative to the NT control (**Fig. 5c-d**). LNPs were then formulated with FLuc mRNA and injected intravenously into gestational day E16 pregnant and nonpregnant mice and luminescence was analyzed 6h later (**Fig. 5e-f, Supplementary Figs. 10-12**). In pregnant mice, NT LNPs delivered luciferase mRNA primarily to the liver and spleen (**Fig. g-h**), whereas all tLNPs showed significantly reduced hepatic and splenic delivery relative to NT LNPs. F(ab’)_2_ tLNPs exhibited significantly enhanced luciferase expression in the placenta compared to NT LNPs, consistent with in vitro performance. Antibody tLNPs demonstrated enhanced placental mRNA delivery compared to PBS treated mice but did not reach statistical significance compared to NT LNPs. None of the formulations showed detectable fetal transfection, consistent with previous reports (**Fig 5i-j**). Overall luminescence in the major organs was higher in nonpregnant mice than pregnant mice (**Fig. 5k-m**). In nonpregnant mice, all formulations localized predominantly to the liver and spleen as expected. Luminescence signals in the liver were comparable across groups, except for antibody tLNPs, which showed reduced hepatic delivery relative to NT LNPs. No significant differences were observed between NT LNPs and tLNPs in the liver or spleen, consistent with the diminished contribution of ligand-mediated selectivity in the absence of EGFR-rich placental tissue.

**Figure 5.**
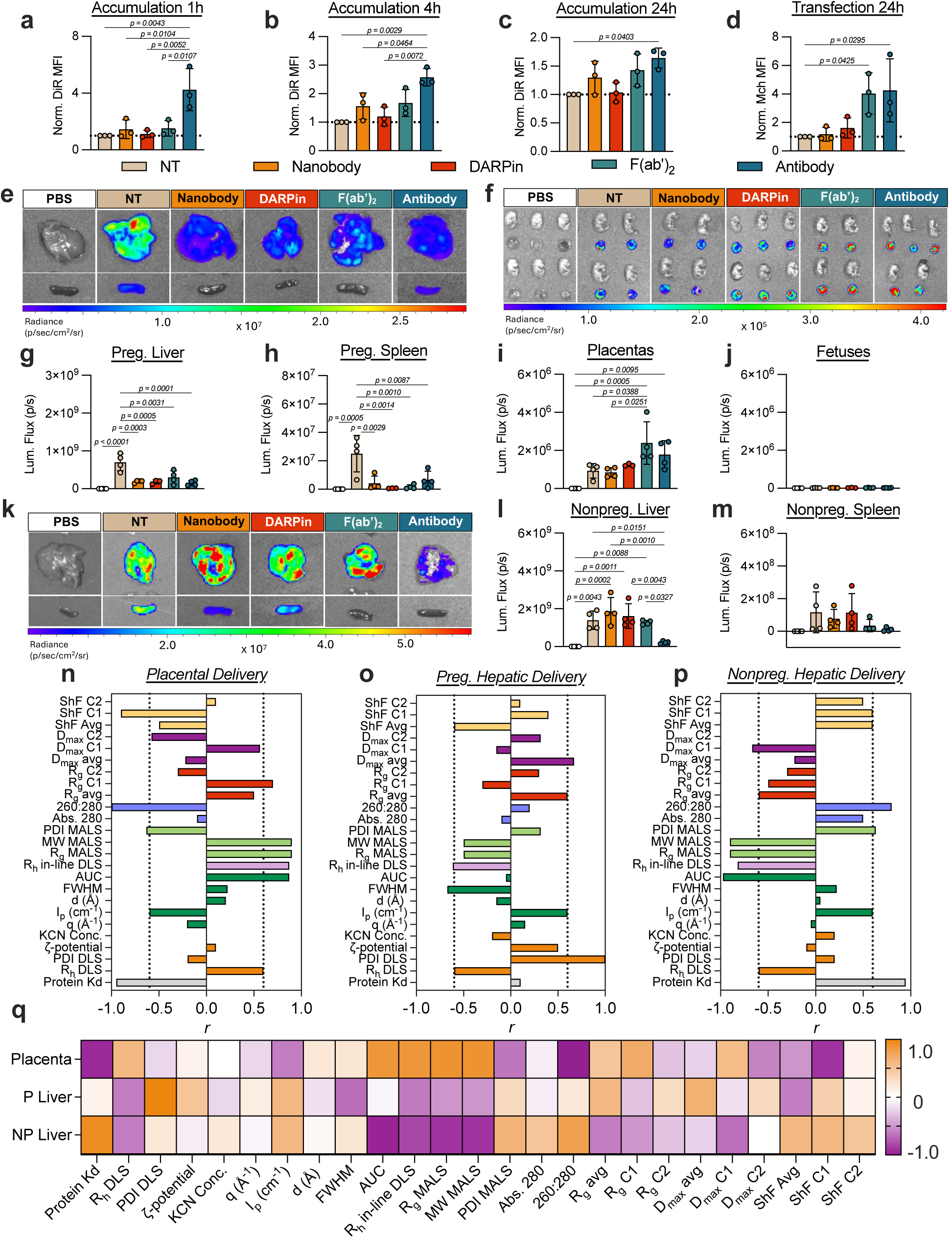
Targeted mRNA delivery to the placenta is driven by tLNP structural subspecies. (**a-d**) DiR-labeled NT LNPs and tLNPs containing mCherry mRNA were incubated with placental BeWo b30 trophoblasts at a dose of 150 ng of mRNA per 150,000 cells. After (**a**) 1 h, (**b**) 4 h, and (**c**) 24 h, cellular accumulation was quantified. After (**d**) 24 h, mCherry expression was also quantified. Normalized DiR and mCherry MFI was calculated by normalizing to cells treated with NT LNPs. (**e–m**) NT LNPs and tLNPs containing FLuc mRNA were administered intravenously via retroorbital injection into pregnant and nonpregnant mice at a dose of 12 µg mRNA per mouse. After 6 h, mice were euthanized, and major organs were dissected. For pregnant mice, luminescence imaging of (**e**) livers and spleens and (**f**) placentas and fetuses were performed via an *in vivo* imaging system (IVIS). Luminescence from (**e-f**) was quantified via region of interest (ROI) analysis to obtain luminescence flux in the (**g**) liver, (**h**) spleen, (**i**) placentas, and (**j**) fetuses of pregnant mice. For nonpregnant mice, luminescence imaging of (**k**) livers and spleens was performed. Luminescence from (**k**) was quantified via region of interest (ROI) analysis to obtain luminescence flux in the (**m**) liver and (**m**) spleen of nonpregnant mice. Signal is reported mean ± SD from *n* = 3 biological replicates for (**a–d**) and *n* = 4 biological replicates for (**e–m).** One-way ANOVA with post hoc Student’s t-tests using the Holm–Sídak correction for multiple comparisons was used to compare fluorescence in for (**a–d**) and luminescence in (**g–h, l–m**) across treatment groups. Nested one-way ANOVA with post hoc Student’s t-tests using the Holm–Sídak correction for multiple comparisons was used to compare luminescence in (**i–j**) across treatment groups. (**n–q**) Spearman correlations for (**n**) placental, (**o**) pregnant hepatic, and (**p**) nonpregnant hepatic luminescence values using the physicochemical parameters from traditional characterization methods, static SAXS analyses, and AF4-UV-DLS-MALS-SAXS analyses. (**q**) Heatmap representing the entire dataset. For Spearman correlation graphs, dotted lines represent *r* = –0.6 and 0.6.

We next correlated measured physicochemical parameters with luminescence outputs using Spearman analysis, defining |r| > 0.6 as a medium-to-strong association (**Fig. 5n-q, Supplementary Table 9**). Placental luminescence showed a strong positive correlation with the area under the curve (AUC) of the batch SAXS Bragg peak, R_h_ determined by in-line DLS, and R_g_ and mass determined by MALS (**Fig. 5n**). In contrast, batch SAXS peak intensity (I_p_), AF4-MALS PDI, and the 260:280 UV-Vis ratio exhibited strong negative correlations. Notably, benchtop protein quantification measurements did not correlate with placental delivery, indicating that simple estimates of total ligand content fail to capture functionally relevant ligand decoration. SAXS-determined R_g_ and D_max_ of C1 correlated positively with placental luminescence, whereas the C1 shape factor correlated negatively, suggesting that a specific, perhaps more isotropic, subpopulation within the tLNP population contributes most strongly to placental targeting. These in vivo correlations were corroborated by in vitro measurements of tLNP binding affinity (**Supplementary Fig. 13a-b, Supplementary Fig. 14**) and inhibition of receptor-mediated endocytosis (**Supplementary Fig. 13c**), followed by biophysical correlation analysis (**Supplementary Fig. 13d-f, Supplementary Table 10**). Consistent with in vivo placental targeting, SAXS-determined R_g_, D_max,_ and the shape factor of C1 showed the strongest correlation with tLNP binding to EGFR-expressing placental cells, and both C1 and C2 parameters correlated with inhibition of tLNP-mediated mRNA delivery upon EGFR blockade.

In pregnant mice, hepatic luminescence correlated positively with standard DLS-derived PDI and batch SAXS peak intensity (I_p_) and negatively with standard DLS-derived R_h_, peak full width half maximum (FWHM) and in-line DLS-derived R_h_ (**Fig. 5o**). SAXS-derived R_g_ and D_max_ of the average profile also correlated positively with hepatic delivery, and the average shape factor correlated negatively, indicating that nonspecific liver delivery is influenced by parameters of the global particle properties rather than by parameters of specific subpopulations. In nonpregnant mice, hepatic luminescence correlated positively with batch SAXS I_p_ and the 260:280 ratio and negatively with batch SAXS AUC, in-line DLS R_h_, and MALS-derived R_g_ and mass (**Fig 5p**). Luminescence also showed negative correlations with SAXS-derived average-population R_g_ and C1 D_max_, and positive correlations with the shape factors of both the average and C1 populations. Collectively, these trends support a coherent model: in pregnant mice, the EGFR-rich placenta enables ligand-mediated targeting, whereas in nonpregnant mice, all LNP subpopulations, including ligand-decorated species, default to hepatic accumulation and subsequent transfection. These findings further suggest that residual nonspecific liver delivery is dictated primarily by bulk physicochemical properties rather than by discrete structural subpopulations (**Fig. 5q**).

### Biophysical-toxicological correlations reveal bulk LNP populations and tLNP subpopulations drive differential acute cytokine responses

We next evaluated whether biophysical parameters were associated with systemic toxicity by measuring serum cytokines (C3a, TNF, IFN-γ, IL-6) and liver enzymes (ALT, AST) in pregnant (**Fig. 6a**) and nonpregnant (**Fig. 6b**) mice. In pregnant mice, nanobody tLNPs elicited significantly elevated C3a levels compared to PBS controls, indicating complement activation. Antibody tLNPs increased TNF and IL-6 and significantly elevated IFN-γ, whereas NT LNPs and DARPin tLNPs produced significantly higher AST levels compared to PBS. Notably, F(ab’)_2_ tLNPs demonstrated the most favorable toxicity profile in pregnant mice, with AST levels significantly lower than NT LNPs and no significant increases in any cytokine observed. In nonpregnant mice, nanobody, DARPin, and F(ab’)_2_ tLNPs produced significantly higher C3a levels compared to PBS. Nanobody, F(ab’)_2_, and antibody tLNPs significantly elevated TNF relative to PBS, with F(ab’)_2_, and antibody tLNPs exceeding NT LNP levels. Antibody tLNPs further induced higher TNF levels compared to F(ab’)_2_ tLNPs. DARPin, F(ab’)_2_, and antibody tLNPs all showed significantly increased IFN-γ levels compared to both PBS and NT LNP treated mice, and antibody tLNPs produced the highest IFN-γ induction among all tLNP formulations. Antibody tLNPs additionally elevated AST relative to both PBS and F(ab’)_2_ tLNPs. No significant differences in ALT were observed in either pregnant or nonpregnant mice.

**Figure 6.**
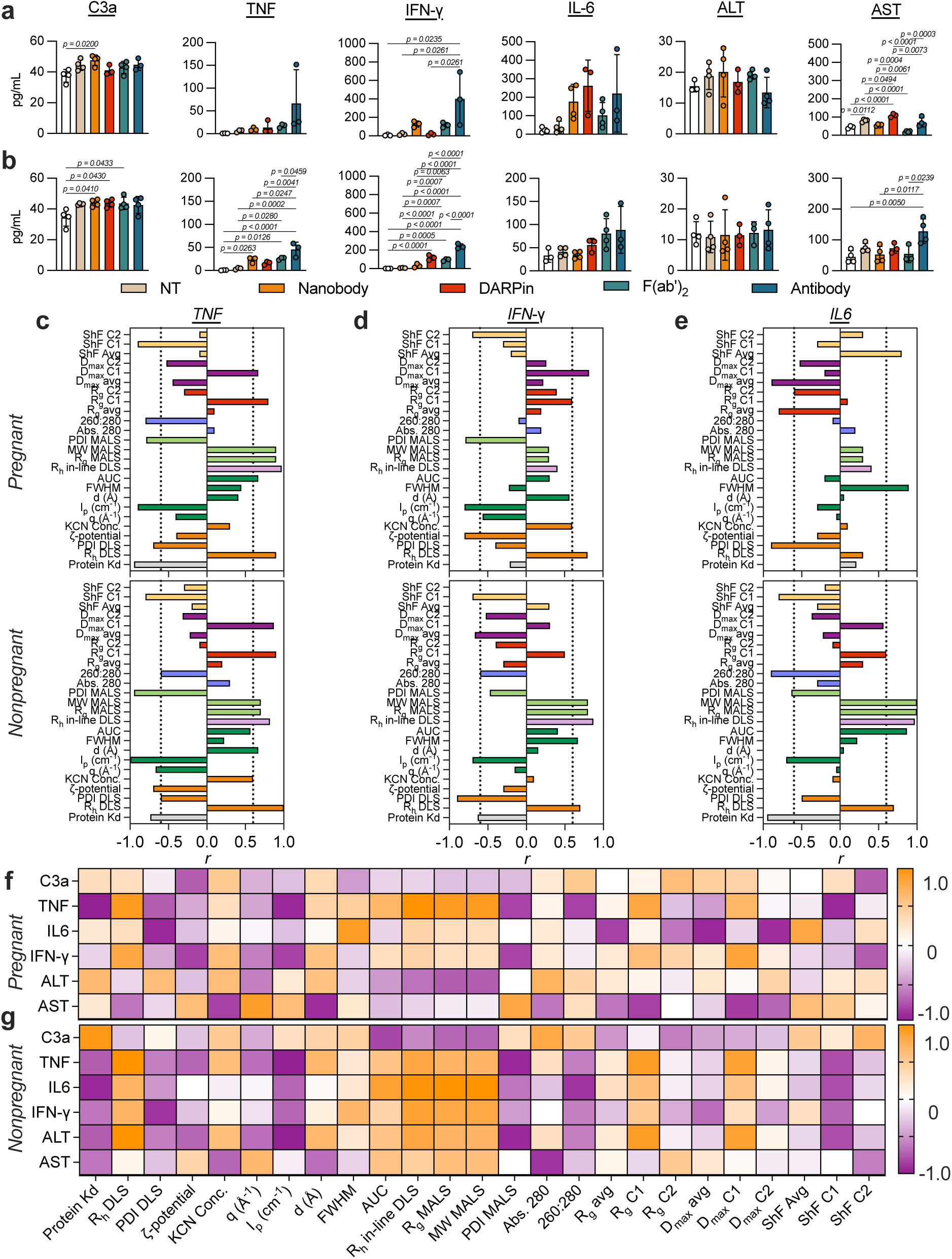
tLNP bulk populations and subspecies drive differential acute cytokine responses. (**a-b**) NT LNPs and tLNPs containing FLuc mRNA were administered intravenously via retroorbital injection into pregnant and nonpregnant mice at a dose of 12 µg mRNA per mouse. After 6 h, mice were euthanized, and serum was collected. Serum levels of C3a, TNF, IFN-γ, IL-6, ALT, and AST were quantified in (**a**) pregnant and (**b**) nonpregnant mice via ELISA. Measurements are reported mean ± SD from *n* = 3–4 biological replicates. One-way ANOVA with post hoc Student’s t-tests using the Holm–Sídak correction for multiple comparisons was used to compare cytokine levels across treatment groups. (**c–e**) Spearman correlations for (**c**) TNF, (**d**) IFN-γ, and (**e**) IL-6 serum levels in pregnant (top) and nonpregnant (bottom) mice using the physicochemical parameters from traditional characterization methods, static SAXS analyses, and AF4-UV-DLS-MALS-SAXS analyses. (**f**) Heatmap representing the entire dataset. For Spearman correlation graphs, dotted lines represent *r* = –0.6 and 0.6.

When the Spearman correlation was applied to cytokine data, several consistent structure–toxicity relationships emerged (**Fig. 6c-g, Supplementary Tables 11-12**). In pregnant mice, cytokine induction (C3a, TNF, and IFN-γ) correlated predominantly with subpopulation-specific features, including positive correlations with SAXS-derived R_g_ and D_max_ of C1 components and negative correlations with their shape factors (**Fig. 6c-d**). Notably, while C1 most strongly correlated with placental luminescence, C3a levels correlated with C2, suggesting the potential existence of a secondary ligand-engaging subpopulation that activates the complement cascade. In addition, TNF correlated negatively with the 260:280 ratio, and TNF, IFN-γ, and AST positively correlated with C1 parameters, suggesting that the structurally defined targeted subpopulation responsible for placental RNA delivery also contributes to acute cytokine induction. In contrast, IL-6 in pregnant mice had strong correlation with average R_g_, D_max_, and shape factor (**Fig. 6e**), which is in agreement with hepatic luminescence correlations, indicating that the classic IL-6 responses often associated with LNP delivery are likely associated with global, ensemble-averaged nanoparticle properties.

The structure-toxicity correlation landscape shifted markedly in nonpregnant mice (**Fig. 6f-g**). TNF again showed positive correlation with C1 parameters and negative correlation with the 260:280 ratio, consistent with LNP-mediated mRNA delivery by the C1 subpopulation. However, IL-6 switched polarity relative to pregnancy, now correlating positively with the R_g_ and D_max_ of C1 rather than with bulk parameters, indicating a switch from global nanoparticle-driven IL-6 production to subpopulation-specific delivery in the liver. C3a and IFN-γ displayed mixed correlations involving both individual component and average profiles, reflecting contributions again from both targeted subpopulations and the averaged population. Features derived from Lorentzian analysis of SAXS profiles and AF4-MALS measurements also contributed to cytokine associations: batch SAXS I_p_ remained negatively correlated with cytokine production across pregnant and nonpregnant mice, FWHM lost its pregnancy-specific positive correlation with IL-6, and both in-line DLS R_h_ and MALS-derived R_g_ and mass showed stronger positive correlations in nonpregnant mice.

## DISCUSSION

Despite rapid advances in targeted RNA delivery, the rational engineering of tLNP systems has remained constrained by the lack of analytical tools capable of resolving the structural heterogeneity that shapes in vivo performance.^20^ In this work, we move beyond ensemble-averaged characterization by integrating AF4-UV-DLS-MALS-SAXS with component-resolved scattering analysis to dissect how ligand attachment reshapes LNP structure, polydispersity, morphology, and functional performance at the subpopulation level, providing enhanced resolution of features that are not accessible by ensemble-averaged methods. This approach enables parallel resolution of compositional heterogeneity (UV chemometrics and light scattering) and structural heterogeneity (SAXS component analysis) across the same AF4-resolved populations.

We find that protein conjugation does not substantially perturb the internal lipid-RNA nanostructure of LNPs but instead remodels their external architecture. Although ligand attachment is generally assumed to leave internal organization intact, this assumption has not previously been directly verified. The preservation of internal order across all tLNPs is consistent with functional differences that arise from changes in external surface composition and geometry rather than from disruption of core assembly. At the same time, ligand attachment dramatically amplifies the intrinsic heterogeneity of LNP formulations, particularly for larger and multivalent proteins. While conventional bulk techniques, such as DLS and SEC-MALS, yield only ensemble-averaged descriptors, AF4-UV-DLS-MALS-SAXS resolves overlapping ligand-dependent tLNP subpopulations with distinct mass, size, and structural features, establishing population-level heterogeneity as a defining consequence of protein functionalization.

A central finding of this work is that ligand decoration remodels LNP geometry, shifting particles from prolate ellipsoid towards increasingly isotropic morphologies as ligand size increases. Leveraging the extended low-q range afforded by our AF4-UV-DLS-MALS-SAXS experimental configuration, we perform Guinier and P(r) analyses that are often challenging for LNPs due to insufficient q-range^54,55^ which enables reliable extraction of R_g_, D_max_, and shape factor for individual fraction-resolved tLNP subpopulations. Analysis of the resulting shape factor (D_max_/R_g_), as defined in the main text, reveals a ligand-size–dependent reduction in particle anisotropy, suggestive of preferential ligand decoration along the elongated axes of the nanoparticle. This hypothesis aligns with the broader concept that LNP elongation may facilitate cell-surface interactions^63–65^ and that changes in LNP morphology impact RNA delivery^66,67^; protein decoration on these axes could reorganize apparent external morphology and enhance target-cell engagement^68^. DENSS reconstructions further support these findings, showing that tLNPs differ from NT LNPs through expansions or contractions along one or more axes, with larger ligands producing greater inter-component variability and, in some cases, more isotropic density envelopes. Our results suggest a mechanistic model in which ligand attachment not only increases particle size but also reshapes external nanoparticle morphology, generating structurally distinct subpopulations, each of which contributes differently to biological targeting and LNP-mediated RNA delivery.

A second, critical finding of this study is that tLNP functional behavior is driven by biophysically defined tLNP subpopulations rather than by bulk-average LNP properties, as revealed by correlations between subpopulation-resolved biophysical parameters and biological outcomes. To our knowledge, this represents the first demonstration that only specific, structurally defined tLNP subpopulations, not ensemble-average properties, correlate with in vivo targeted mRNA delivery. Specifically, targeted placental mRNA delivery correlates most strongly with the R_g_, D_max_, and shape factor of C1 components, whereas ensemble-averaged parameters instead predict nonspecific hepatic transfection. Cytokine induction follows a similar pattern in pregnant mice: TNF and IFN-γ track with C1 population features, whereas IL-6 correlates with ensemble-averaged values. When the placental target is absent, mRNA delivery and cytokine correlations shift toward a mixture of individual components and bulk metrics, reflecting simultaneous contributions from ligand-decorated populations and the global LNP population. These results indicate that targeted delivery is likely mediated primarily by one or more structurally and compositionally defined subpopulations rather than by the entire heterogeneous ensemble. This interpretation is consistent with the intrinsic inefficiency of LNP formulations, in which only a fraction of LNPs encapsulate RNA^19,69^, and only a subset of those are successfully decorated with targeting ligands^70^. Therefore, the biologically active tLNP species (those containing both RNA and protein ligand) are likely a small subset within the bulk formulation. These findings point to a clear next step for the field: the fractionation and isolation of discrete tLNP subpopulations to directly interrogate their functional contributions. Enriching or purifying these subpopulations may significantly improve therapeutic performance and reduce off-target effects, marking a conceptual shift from bulk formulation optimization toward subpopulation-level nanoparticle engineering.

An important implication of this subpopulation-driven framework is that increased structural heterogeneity does not necessarily preclude effective targeting, as exemplified by the superior performance of F(ab’)_2_ tLNPs. In pregnant mice, F(ab’)_2_ tLNPs result in the highest placental luminescence with minimal toxicity. This disparity likely reflects the unique advantages of avidity: F(ab’)_2_ and antibody ligands possess dual binding sites and lower K_d_ values than monovalent nanobody and DARPin ligands. Avidity is multiplicative rather than additive, and may partially compensate for structural heterogeneity by increasing the probability of receptor engagement and subsequent endocytosis^71^. This principle is consistent with principles observed in biological systems such as viral capsids, which rely on probabilistic multivalent interactions to overcome stochastic barriers during nuclear entry^72^. Our findings therefore suggest that ligand avidity may be a key mechanism enabling tLNPs to function effectively despite their pronounced polydispersity.

Collectively, this work is, to our knowledge, the first to reveal that the functional behavior of tLNPs is not governed by ensemble-averaged properties but instead by distinct, structural subpopulations that have historically remained analytically inaccessible. While separation-coupled, solution-phase biophysical platforms have been applied to mRNA LNPs, here we extend this integrated approach to tLNP systems, enabling the first direct resolution of functionally relevant tLNP subpopulations and, further, the first correlation of their nanoscale structure to in vivo targeting performance. We anticipate that the AF4-UV-DLS-MALS-SAXS platform and subsequent chemometric analyses used in this work will inform a mechanistic, data-driven framework to guide future improvements in rational tLNP engineering. Importantly, this framework is developed and validated in the context of pregnancy, a uniquely stringent physiological setting in which therapeutic choices for pregnancy complications remain scare and safety constraints render systemic delivery approaches poorly suited^16,32,33^. Under these conditions, therapeutic efficacy is unlikely to scale with bulk-average biodistribution and will demand tLNP engineering at the subpopulation level to achieve exceptional targeting precision. More broadly, this work addresses a critical gap in nanotechnology and drug delivery research, where mechanistic structure–activity relationships for diseases and therapeutic contexts unique to women’s health have remained largely underexplored. By resolving tLNP heterogeneity at the subpopulation level and linking defined structural features to in vivo outcomes, this work enables the generation of high-quality, interpretable datasets that move beyond bulk descriptors. Importantly, such datasets are well suited for integration with machine-learning and AI-driven optimization strategies, which will enable the derivation of predictive design rules for targeted delivery systems. Together, these findings establish that identifying, enriching, and engineering functionally competent tLNP subpopulations will be essential for the development of next-generation precision tLNP-mediated RNA therapeutics not only in pregnancy and women’s health, but also other therapeutic settings where targeting accuracy is non-negotiable.

## METHODS

### Materials

Cholesterol was purchased from Sigma Aldrich. The remaining lipid excipients were purchased from Avanti Polar Lipids. Anti-human EGFR antibody (Cetuximab biosimilar) was purchased from Bioxcell. FLuc and mCherry mRNA with N^1^-methylpseudouridine modifications was purchased from TriLink Biotechnologies. DiR, 10X PBS (pH 7.4), enzymatic digestion kits, EDTA (pH 8.0), Sytox Blue, and dialysis cassettes were purchased from Thermo Fisher Scientific. Citrate buffer was purchased from Teknova. Dulbecco’s Modified Eagle Medium (DMEM), Opti-MEM Reduced Serum Medium, fetal bovine serum (FBS), and penicillin-streptomycin (P/S) were purchased from Gibco. 96-well and 48-well plates were purchased from Corning. Dynasore, bafilomycin A1, amiloride, wortmannin, cytochalasin D, and methyl-beta-cyclodextrin small molecule inhibitors were purchased from Sigma Aldrich. Erlotinib small molecule inhibitor was purchased from Cayman Chemical. The Luciferase Reporter 1000 Assay System and CellTiter-Glo Luminescent Assay System were purchased from Promega. D-Luciferin Potassium Salt was purchased from Regis Technologies.

### Protein modification

Anti-EGFR nanobodies (9G8)^73^ and DARPins (E01)^74^ were expressed and purified using the previously described sortase-tag expressed protein ligation system (STEPL)^75^. To generate expression plasmids, gBlocks for either 9G8 or E01 were inserted into a pSTEPL backbone between NdeI and AgeI restriction sites using In-Fusion Cloning (Takara Bio USA). Proteins were expressed in T7 Express Competent E. coli (C2566, New England Biolabs) and purified with a single DBCO handle attached to the C-terminal via STEPL. Proteins were further purified via size exclusion chromatography on a Superdex^TM^ 75 Increase 10/300 GL Column (Cytiva). Protein concentration was determined via BCA assay. The purified nanobody and DARPin proteins were stored at -20°C for later use.

Anti-EGFR cetuximab antibody was cleaved into F(ab’)_2_ fragments using a ficin enzymatic digestion kit and purified using a Protein A column (Thermo Fisher Scientific). Whole antibodies and F(ab’)_2_ fragments were functionalized with DBCO via a reaction with a 20-fold molar excess of TFP-PEG(4)-DBCO (Thermo Fisher Scientific) in anhydrous DMSO for 2 h at 25°C. Unreacted TFP-PEG(4)-DBCO was removed using Zeba Dye and Biotin Removal spin columns (Thermo Fisher Scientific). Final protein concentration was measured using a Qubit Protein Quantification Assay (Thermo Fisher Scientific). The purified DBCO-labeled antibodies were stored at 4°C for later use.

### tLNP formulation

tLNPs containing lipid A4 were formulated as previously described^38^. Lipid excipients were dissolved in ethanol to form the organic phase using the following molar ratio: ionizable lipid A4 (35 mol.%), DOPE (16 mol.%), cholesterol (46.5 mol.%), C14-PEG2000 (2.1 mol.%), and DSPE-PEG2000-azide (0.4 mol.%). mRNA was dissolved in citrate buffer (pH 3) to form the aqueous phase. LNPs were formulated at a weight ratio 10:1 of ionizable lipid to RNA with a volume ratio of organic phase to aqueous phase of 1:3. The organic and aqueous phases were loaded into glass syringes (Hamilton) and attached to a Pump 33 DDS syringe pump (Harvard Apparatus). The two phases were pushed through a custom soft lithography-patterned polydimethylsiloxane (PDMS) herringbone microfluidic chip, fabricated as described in our previous study^35^. The organic and aqueous phases were injected at a flow rate of 0.6 ml min^−1^ and 1.8 ml min^−1^, respectively. Empty LNPs were prepared in the same manner, but with an aqueous phase containing citrate buffer and no mRNA. For in vitro flow cytometry studies, LNPs were labeled with 1.0 mol.% DiR before loading into dialysis cassettes. For remaining studies, LNPs were loaded directly into dialysis cassettes. LNPs were dialyzed against 1X PBS (pH 7.4) for 2 h in cassettes with a 20kDa molecular weight cutoff filter (Thermo Fisher Scientific). Non-targeted LNPs were stored at 4°C for later use.

To functionalize LNPs with protein, DBCO-labeled proteins were added to LNPs at a 1:200 ratio of DBCO-protein to lipid-anchored PEG-azide and incubated for 18 h at 25°C with gentle shaking. tLNPs were purified from residual, unbound protein using size exclusion chromatography. Briefly, a column was packed with Sepharose CL-6B (Sigma Aldrich) and rinsed with 1X PBS (pH 7.4) to clear ethanol from the system. tLNPs were passed through the column and collected in a 96-well plate. All fractions containing mRNA were pooled and concentrated using 100kDa filters (Sigma Aldrich). The final tLNP solution was stored at 4°C for later use.

### DLS

LNP R_h_, PDI, and particle concentration were determined via dynamic light scattering (DLS) measurements with a DynaPro Plate Reader III (Wyatt Technology) using a cumulants model. LNPs were diluted 15-fold in 1X PBS (pH 7.4) and loaded into a 384-well Aurora plate (Wyatt Technology). The plate was centrifuged at 300*g* for 1 min and then loaded onto the plate reader. Size is reported as intensity-weighted averages from *n* = 3 independent measurements per sample and data are expressed as the mean ± SD, where SD was calculated with the following formula: 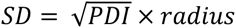

### ζ-potential

ζ-potential was determined via electrophoretic light scattering (ELS) measurements on a Zetasizer Nano (Malvern Instruments). LNPs were diluted 50-fold in ultrapure water and then transferred to disposable folded capillary cells (Malvern Instruments). For each sample, *n* = 3 measurements with *n* = 5 runs were recorded.

### Fluorescent protein quantification

Protein content of tLNPs was estimated using the CBQCA Protein Quantitation Kit according to the manufacturer’s instructions (Thermo Fisher Scientific). BSA standards and LNP samples were prepared in 0.1 M sodium borate containing 0.1% (v/v) Triton X-100 and then transferred to a black 96-well plate. 20 mM KCN and 40 mM ATTO-TAG CBQCA reagent were added to the standards and samples to a total reaction volume of 100 μL at the ratios specified in the manufacturer’s instructions. The plate was shaken at 300 rpm for 1 h at 25°C and fluorescence was read at an excitation wavelength of 465 nm and an emission wavelength of 550 nm. LNP protein content was estimated by first subtracting the base fluorescence from an LNP with no protein and then comparing with a standard curve fitted using univariate least-squares linear regression.

### LNP encapsulation

Relative encapsulation efficiency and encapsulated RNA concentration were determined using a Quant-iT RiboGreen assay (Thermo Fisher Scientific). Each LNP formulation was diluted 80X in either 1X tris-EDTA (TE) buffer (Thermo Fisher Scientific) or 1X TE buffer containing 0.1% (v/v) Triton X-100 (Sigma Aldrich). LNPs were shaken at 300 rpm for 10 min at 25°C to facilitate particle lysis and then transferred to a black 96-well plate. A standard curve was prepared according to the manufacturer’s instructions. RiboGreen fluorescent detection reagent was added to each well at a 1:1 (v/v) ratio per the manufacturer’s instructions. After incubation for 5 min at 25°C, the fluorescence intensity was read on an Infinite 200 Pro plate reader (Tecan) at an excitation wavelength of 480 nm and an emission wavelength of 520 nm. Encapsulation efficiency was calculated as (B-A)/B X 100%, where A is the measured RNA content in TE buffer and B is the measured RNA content in Triton X-100. mRNA concentrations of LNP formulations were estimated by comparison with a standard curve fitted using univariate least-squares linear regression.

### Size-exclusion chromatography in-line with multiangle light scattering (SEC-MALS)

SEC-MALS experiments were performed on 100 µL injections of ∼10^12^ particles/mL samples at 0.5 ml/min at room temperature in 1x PBS buffer using a TOSOH TSKgel G6000PWXL-CP SEC column (7.8 mm × 300 mm, 13-μm particle size; Millipore Sigma). Absolute molar mass was determined in-line using multi-angle light scattering from the column eluent at 18 angles using a DAWN-HELEOS II MALS detector (Waters/Wyatt Technology Corp.) operating at 658 nm. Eluent concentration was recorded using an in-line Optilab T-rEX Interferometric Refractometer (Waters/Wyatt Technology Corp.). The online MALS detectors were normalized for counting efficiency using BSA monomer peak data from separate experiment on a Superose 6 10/300 column in the same running buffer.

For compositional analysis of the base nontargeted mRNA-LNP, data acquisition and analysis were performed as previously described^24^ using the LNP Analysis Module in the ASTRA^TM^ 8.2.2 software (Waters/Wyatt Technology Corp.). To obtain accurate concentration data from the online 260 nm UV absorbance signal, empty LNP samples were also prepared using the microfluidic devices with lipid compositions and concentrations matching the LNP-mRNA samples. The empty LNP samples were measured with the same method as the LNP-mRNA samples, and the results from the empty samples were used to generate experimentally derived UV scattering correction profiles with the ASTRA LNP Analysis Module as previously described^41^. The scattering correction profiles were then applied non-destructively to the data collected for the corresponding LNP-mRNA samples.

For mass determination of protein-labeled mRNA LNP particles, the weight-averaged molar masses of species within defined chromatographic peaks were calculated using the ASTRA software version 8.2.2 (Waters/Wyatt Technology Corp.), by construction of Debye plots (KC/Rθ versus sin^2^[θ/2]) at one second data intervals. The weight-averaged molar mass was then calculated at each point of the chromatographic trace from the Debye plot intercept, and an overall average molar mass was calculated by weighted averaging across the peak. A mass-averaged *dn/dc* for the composite particle was calculated using the compositional analyses of the base mRNA-LNP performed and the calculated number of surface azide attachment sites per particle, assuming 100% labelling efficiency, yielding calculated *dn/dc* figures ranging from 0.160–0.165 mL/g. These derived properties are provided in **Supplementary Table 2**.

### Synchrotron batch small angle X-ray scattering (SAXS)

Batch SAXS data were collected at beamline 16-ID (LiX) of the National Synchrotron Light Source II (Upton, NY)^76^. SAXS/WAXS data were simultaneously collected at a wavelength of 0.89 Å, yielded accessible scattering angle where 0.006 < q < 3.0 Å^-1^, where q is the momentum transfer, defined as 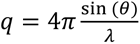, where *λ* is the X-ray wavelength and *2θ* is the scattering angle; data to q<0.5 Å^-1^ were used in subsequent analyses. 60 μL of sample at a particle concentration of ∼10^12^ particles / mL were examined. Samples in dialysis matched 1x PBS (pH 7.4) were loaded into a 1-mm capillary for ten 1-s X-ray exposures.

### Sedimentation velocity analytical ultracentrifugation (SV-AUC)

Sedimentation velocity analytical ultracentrifugation experiments were performed at 20°C with an Optima^TM^ analytical ultracentrifuge (Beckman-Coulter) and a TiAn50 rotor with two-channel charcoal-filled Epon centerpieces and sapphire windows, using both absorbance and interference optics. Data were collected in 1x PBS with detection at 260 & 280 nm, as well as interference optics. Complete sedimentation velocity profiles were recorded every 30 seconds at 20,000 rpm and 20°C. Data were fit using a least-squares *g*(s)* model as implemented in the program SEDFIT^77^.

### Synchrotron AF4-UV-DLS-MALS-SAXS

Data were collected at the BioCAT beamline (18ID) at the Advanced Photon Source at Argonne National Labs (Argonne, IL)^78^. SAXS data were simultaneously collected at a wavelength of 1.033 Å (12.0 keV), yielded accessible scattering angle where 0.0025 < q < 0.413 Å^-1^, where q is the momentum transfer, defined as 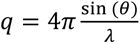,where λ is the X-ray wavelength and 2θ is the scattering angle; data to q < 0.2 Å^-1^ were used in subsequent analyses. 100 μL of LNP at a particle concentration of ∼10^12^ particles were injected and eluted isocratically from an Eclipse^TM^ NEON field flow fractionation (AF4) instrument with dilution control module using a variable-height short channel with a 275 µm spacer and a 10 kDa regenerated cellulose membrane (Wyatt Technology), connected to a 1260 Infinity II HPLC system with a G711A quaternary pump and a G7167A multisampler (Agilent Technologies). A DAWN^TM^ HELEOS II MALS instrument with an integrated WyattQELS^TM^ DLS detector (Wyatt Technology), an Optilab^TM^ T-rEX differential refractometer (Wyatt Technology), and a G7165A UV-Vis multi-wavelength UV-Vis detector (Agilent Technologies) were used for online detection. 1X PBS (pH 7.4) was used as the mobile phase. Sample injection volumes were 55–150 μL. The AF4 system was controlled by VISION 3.2.0 software (Wyatt Technology). AF4 method parameters: 2.5 mL min^-1^ channel flow and 0.5 mL min^-1^ detector flow (corresponding to a 5-fold concentration enhancement from the dilution control module), 0.2 mL min^-1^ inject flow. Focusing was at 1.0 mL min^-1^ with a 25% focus position for 14 min. Cross flow was ramped from 0 to 1.0 mL min^-1^ in the first minute of the method, and then kept constant at 1.0 mL min^-1^ throughout focusing and the initial 5 min of the elution step, followed by an exponential gradient of 1.0–0.04 mL min^-1^ over 30 min after which the crossflow was kept constant at 0.04 mL min^-1^ for 10 min before being turned off. Eluent from the AF4, after flowing through the UV, MALS-DLS and RI detectors, was flown through the SAXS flow cell. The flow cell consists of a 1.0 mm ID quartz capillary with ∼20 µm walls. A coflowing buffer sheath is used to separate sample from the capillary walls, helping prevent radiation damage^79^. Scattering intensity was recorded using an EIGER2 XE 9M (Dectris) detector which was placed 3.714 m from the sample using0.15 s X-ray exposures taken at 1 second intervals over the entire elution and data was reduced to 1D scattering profiles using BioXTAS RAW 2.3.0^80^. Plots of intensity from the forward scatter closely correlated to in-line UV and refractive index (RI) measurements.

#### In-line Absorbance Analyses

In-line diode-array spectral data was collected at eight wavelengths (4 nm bandpass) between 210-300 nm across 3634 time points. Chemical spectral and concentration profiles were resolved using Multivariate Curve Resolution – Alternating Least Squares (MCR-ALS) implemented via the pyMCR library^81^. All chemometric analyses were performed in Python 3.10 using the Anaconda distribution on Ubuntu Linux, using *NumPy*, *pandas*, *SciPy*, *scikit-learn*, *pyMCR*, and *matplotlib*. Data handling used *NumPy* and *pandas*. Prior to multivariate analysis, chromatographic matrices were converted into sample × wavelength and wavelength × sample representations. No baseline subtraction or smoothing was applied unless specified. To determine the intrinsic dimensionality of each chromatographic dataset, singular value decomposition (SVD) was applied^50^. Singular values were normalized to the first singular value, and components with normalized singular values above 0.03 were retained as significant. The resulting SVD rank serves as an upper bound on the number of chemically meaningful species. To refine the chemical rank, Evolving Factor Analysis (EFA)^51^ was applied in both forward and backward directions. At each step, SVD was performed on growing (forward) or shrinking (backward) subsets of the chromatogram. A component was considered active if its normalized evolving singular value exceeded 0.03 at any point. The final chemical rank was defined as the minimum of SVD rank, forward EFA rank, and backward EFA rank. PCA^51,82^ was used to obtain initial spectral estimates, with the number of components equal to chemical rank plus one (to model baseline contributions). Alternating least-squares iterations were solved under non-negativity constraints on both concentration and spectral profiles. The first MCR component was assigned to protein absorbance and the second to RNA, based on UV–Vis characteristics. Normalized concentration profiles were computed, and RNA-to-protein ratios were determined pointwise to assess component co-elution and relative abundance across the chromatographic time domain. Gaussian models containing one, two, or three components were fit to the major protein elution peak using nonlinear least-squares optimization. Model selection was performed using the Akaike Information Criterion (AIC), with the lowest AIC indicating the best model. Residuals and fit diagnostics were recorded for all candidate models.

#### In-line MALS/DLS Analyses

Prior to LNP sample injections in the AF4-MALS component, the membrane was conditioned by injecting 100 μL of a 4 mg mL^-1^ bovine serum albumin (BSA Fraction V) standard (Wyatt Technology). The system performance was checked by injecting and analyzing a triplicate of 25 μL of the same BSA standard using the BSA short channel AF4 method built into the VISION 3.2.0 software. The online MALS detectors were calibrated at 90° scattering angle and the remaining detector angles normalized to the response at 90° using the BSA monomer peak data. Data acquisition and analysis were performed using ASTRA^TM^ 8.2.1 software (Wyatt Technology). Masses were calculated using the mass-averaged *dn/dc* values in **Supplementary Table 2**.

#### SAXS analyses

Analysis of the AF4-SAXS datasets was performed using the BioXTAS RAW program^80,83^. Buffer-subtracted profiles were analyzed by SVD and the ranges of overlapping peak data were determined using EFA as implemented in REGALS^52,84^. The determined peak windows were used to identify the basis vectors for each component and the corresponding SAXS profiles were calculated. When manually fitting the pair distribution function P(r), the maximum diameter of the particle (D_max_) was incrementally adjusted in GNOM^85^ as implemented in RAW to maximize the total estimate and χ^2^ figures, to minimize the discrepancy between the fit and the experimental data to a q_max_ of 0.1 Å^−1^ and to optimize the visual qualities of the distribution profile.

Deconvolution of the primary Bragg peaks in SAXS were using multiple Lorentz fits^45^ as implemented in the program Origin (OriginLab Corporation):

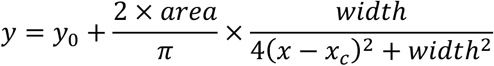

where y_0_ is an offset and was set to the X-ray baseline. x_c_ is a center of function and corresponds to the center of the SAXS peak.

The high-q region of each SAXS profile was analyzed using Porod scaling to probe short-length-scale density correlations. In the Porod regime, the scattered intensity follows a power-law dependence, 𝐼(𝑞) ∼ 𝑞^,-^, where the Porod exponent 𝑚 reflects interfacial and internal density fluctuations at length scales smaller than the overall particle size. Ideal compact particles with sharp interfaces exhibit classical Porod behavior (𝑚 = 4), whereas deviations toward lower values indicate increasing structural heterogeneity. Exponents in the range 3 < 𝑚 < 4 are associated with surface-fractal scattering arising from rough or diffuse interfaces, while values 1 < 𝑚 < 3 correspond to mass-fractal scattering from internally disordered structures^86–88^. Porod exponents were determined by linear fitting of log 𝐼(𝑞) versus log 𝑞 over the experimentally accessible high-q range using ScÅtter^89^. Fits were applied consistently across all datasets, including batch measurements, elution-resolved, and component-resolved profiles obtained by singular value decomposition (SVD). The resulting exponents were used to classify particle structural behavior without assuming idealized geometries.

DENSS^90^ was used to calculate the ab initio electron density map directly from the GNOM output. In total, 20 reconstructions of electron density were performed in slow mode with default parameters and subsequently averaged and refined. Reconstructions were visualized using the PyMOL 2.5.2 Molecular Graphics System (Schrödinger) with five contour levels of density rendered with the following colors: 15σ, red; 10σ, green; 5σ, cyan; 2.5σ, blue; −0.7σ, blue. To determine size parameters, DENSS density maps were converted to pseudo-atomic PDB format using the vol2pdb utility from the Situs package^59^.

### Cell culture

A431 cells were kindly provided by Dr. Andrew Tsourkas (University of Pennsylvania). BeWo b30 cells were kindly provided by Dr. Dogeun Huh (University of Pennsylvania) with permission from Dr. Alan Schwartz (Washington University in St. Louis). BeWo b30 cells and A431 cells were cultured in Dulbecco’s Modified Eagle Medium (DMEM) supplemented with 10% fetal bovine serum (FBS) and 1% penicillin-streptomycin (P/S) and maintained at 37°C and 5% CO_2_.

### In vitro transfection studies

For luminescence assays, A431 cells were plated at a density of 25,000 cells/well in 100 µL DMEM in tissue-culture treated 96-well plates and then left to adhere overnight. DMEM was aspirated and replaced with Opti-MEM Reduced Serum Medium. Cells were then treated with LNPs encapsulating FLuc mRNA at a dose of 25 ng of mRNA per well. After 24 h, media was removed, and cells were incubated with 0.1% Triton-X for 3 min. 100 μL of luciferase assay substrate (Promega) was then added to each well, and cells were left to incubate at room temperature for 5 min. Luminescence was detected using an Infinite 200 Pro plate reader (Tecan). Normalized luciferase expression for each tLNP treatment group was quantified by normalizing to cells treated with non-targeted LNPs.

For flow cytometry studies, BeWo b30 cells were plated at a density of 150,000 cells/well in 200 µL DMEM in tissue-culture treated 48-well plates and then left to adhere overnight. DMEM was aspirated and replaced with Opti-MEM Reduced Serum Medium. Cells were then treated with DiR-labeled LNPs encapsulating mCherry mRNA at a dose of 150 ng of mRNA per well. After 1 h, 4 h, or 24 h, media was removed, and cells were washed with 1X PBS (pH 7.4) containing 2 mM EDTA. Cells were then resuspended 1X PBS-EDTA containing 0.125% (v/v) Sytox Blue. Single cell suspensions were analyzed via flow cytometry for DiR and mCherry fluorescence using a BD LSR II flow cytometer equipped with violet, blue, green and red lasers. For each sample, at least 10,000 events within the singlet gate were collected. Data analysis was performed using FlowJo version 10.9 (Becton Dickinson). Normalized DiR or mCherry median fluorescent intensity (MFI) for each treatment group was quantified by normalizing to cells treated with non-targeted LNPs at each time point.

### In vitro endocytosis studies

BeWo b30 cells were plated at a density of 25,000 cells/well in 100 µL DMEM in tissue-culture treated 96-well plates and then left to adhere overnight. Cells were treated with small molecule endocytosis inhibitors dissolved in DMSO for 30 min. at the following concentrations: dynasore (100 µM), amiloride (2 mM), methyl-beta-cyclodextrin (5 mM), and erlotinib (200 µM). Cell media was then aspirated and replaced with Opti-MEM Reduced Serum Medium. Cells were then treated with LNPs encapsulating FLuc mRNA at a dose of 25 ng of mRNA per well. Control cells were treated with DMSO alone (no inhibitor) followed by LNPs. After 24 h, luciferase expression was evaluated using a Luciferase Reporter 1000 Assay System. Cell viability was evaluated using a CellTiter-Glo Luminescent Assay System. To calculate toxicity, each group was normalized to BeWo b30 cells treated with DMSO containing no inhibitor followed by treatment with LNPs. To calculate relative luminescence, each group was first normalized to BeWo b30 cells treated with DMSO containing no inhibitor and then normalized to each group’s estimated toxicity value to account for potential decreases in luminescence due to DMSO-mediated toxicity.

### Animal experiments

All animal use was in accordance with the guidelines and approval from the University of Pennsylvania’s Institutional Animal Care and Use Committee (IACUC, protocol #806540). Female and male C57BL/6 mice (8-12 weeks old, approximately 20 g average weight) were purchased from the Jackson Laboratory. Time-dated pregnancies were achieved by housing two females and one male all at least 8–12 weeks of age together in one cage overnight and separating the male the next morning. Consistent with the breeding scheme established by Jackson Laboratory, separation was deemed to be gestational day E0. Mice were housed in a vivarium with a 12 h light–dark cycle and were provided with food and water ad libitum. Vivarium temperature was maintained between 20 and 24 °C and humidity was maintained between 40 and 70%.

### In vivo biodistribution studies

LNPs encapsulating FLuc mRNA were administered to non-pregnant and gestational day E16 pregnant mice via retroorbital injection at a dose of 12 µg mRNA per mouse. After 6 h, mice received an intraperitoneal injection of *D-*luciferin potassium salt at a dose of 150 mg kg^-1^. After 5 min, blood was collected by retroorbital bleeding in Microtainer blood collection tubes containing serum separator gel (BD Biosciences). Mice were then euthanized with CO_2_ and the heart, lung, liver, kidneys, spleen, and uterus were removed. In pregnant mice, the uteruses were dissected to remove the fetuses and placentas. Luminescence imaging of the organs was performed using an in vivo imaging system (IVIS; PerkinElmer). Luminescence flux was quantified using the Living Image Software 4.7.3 (PerkinElmer) by placing rectangular regions of interest (ROIs) on organ images, keeping the same ROI sizes among each organ. Collected blood was allowed to clot for 2 h at room temperature and then centrifuged for 20 min at 2000 g. Serum was removed, aliquoted, and stored at -20 °C for later use. Mouse Quantikine ELISA kits (R&D systems) were used to evaluate C3a, IL-6, TNF, and IFN-γ levels in serum per the manufacturer’s instructions. Colorimetric assay kits (Cayman Chemical) were used to evaluate ALT and AST levels in serum per the manufacturer’s instructions.

### Statistical analysis

All statistical analysis was performed in GraphPad Prism version 10.6.0. All tests of significance were performed at a significance level of α = 0.05. Ordinary (one technical replicate per biological replicate) or nested (multiple technical replicates per biological replicate) one-way ANOVAs with post hoc Student’s t-tests using the Holm–Šídák correction for multiple comparisons were used to compare responses across treatment groups. Each fetus or placenta from one mouse was considered a technical replicate. Two-sided ordinary two-way ANOVAs with post hoc Student’s t-tests using the Holm–Šídák correction for multiple comparisons were used to compare responses across treatment groups and gestational days or mouse cohorts (nonpregnant versus pregnant). All data are presented as the mean ± standard deviation (SD) with the exact number of replicates reported in the figure legends. All samples had at least *n* = 3 biological replicates, and no statistical methods were used to predetermine sample size. Investigators were not blinded during outcome assessment or data analysis.

## DATA AVAILABILITY

All data supporting the findings of this study are available within the paper and Supplementary Information. The SEC-MALS, SAXS, and AF4-UV-DLS-MALS-SAXS raw and analyzed data are available from Zenodo. Source data are provided with this paper.

## ACKNOWLEDGEMENTS

The authors acknowledge and thank the Penn Cytomics and Cell Sorting Core (RRID: SCR_022376). The LiX beamline is part of the Center for Biomolecular Structure (CBMS), which is supported by the Department of Energy Office of Biological and Environmental Research (KP1605010). As part of NSLS-II, a national user facility at Brookhaven National Laboratory, work performed at the CBMS is supported in part by the US Department of Energy, Office of Science, Office of Basic Energy Sciences Program under contract number DE-SC0012704. This research was performed on an APS beam time award from the Advanced Photon Source, a U.S. Department of Energy (DOE) Office of Science user facility operated for the DOE Office of Science by Argonne National Laboratory under Contract No. DE-AC02-06CH11357. BioCAT was supported by grant P30 GM138395 from the National Institute of General Medical Sciences of the National Institutes of Health. K.G. acknowledges support from the Johnson Research Foundation.

M.J.M. acknowledges support from an NSF CAREER Award (no. CBET-2145491) and a Burroughs Wellcome Fund Career Award at the Scientific Interface. H.C.G, H.C.S., A.S.T., H.M.Y, and A.A. were supported by National Science Foundation Graduate Research Fellowships (Award 1845298). The content is solely the responsibility of the authors and does not necessarily reflect the official views of the National Institute of General Medical Sciences or the National Institutes of Health.

## AUTHOR CONTRIBUTIONS

H.C.G., M.S.P., M.B.W., J.B.H., and K.G. conceived and designed the experiments. H.C.G., H.C.S., A.S.T., M.S.P., E.B., H.M.Y., V.M.U., A.C., B.E.N., A.A., M.B.W., J.B.H., and K.G.

performed the experiments. H.C.G., M.B.W., J.B.H., and K.G. analyzed the data. H.C.G., K.G., and M.J.M wrote and edited the manuscript. A.T., K.G., and M.J.M. funded the experiments. All authors discussed the results and approved the final version of the manuscript for submission.

