## Supplementary Information for "Resolving heterogeneity of targeted lipid nanoparticles through solution-based biophysical analyses"

*Michael J. Mitchell	*Kushol Gupta

Associate Professor	Research Assistant Professor

Department of Bioengineering	Department of Biochemistry and Biophysics

University of Pennsylvania	University of Pennsylvania

25 N 38^th^ St	422 Curie Blvd.

Suite 1014	901C Stellar-Chance Building

Philadelphia, PA 19104, USA	Philadelphia, PA 19104, USA

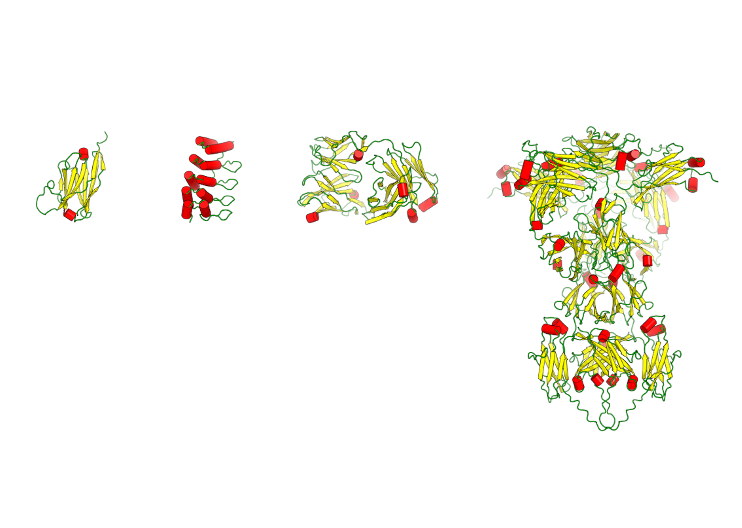
**SUPPLEMENTARY FIGURES**

**Supplementary Figure 1 – Predicted structures of proteins used to generate tLNPs.** From left to right: nanobody, designed ankyrin repeat protein (DARPin), F(ab’)_2_ fragments, and whole antibody. Models were generated using Alphafold3^1^. Yellow denotes predicted b-sheets, red denotes alpha helices, and green represents regions with no predicted secondary structure.

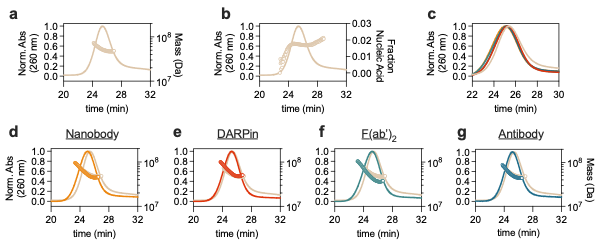

**Supplementary Figure 2 – NT LNPs and tLNPs characterized via SEC-MALS.** (**a**) Molar mass profile of NT LNPs determined by SEC-MALS. (**b**) Conjugate analysis enabled determination of nucleic acid content, confirming the expected two-component lipid-RNA composition of NT LNPs. (**c**) Size-exclusion profiles overlaid of NT LNPs with all tLNPs. All tLNPs eluted earlier than NT LNPs, suggesting increased size of tLNPs. Absorbance at 260 nm and molar mass profiles of (**d**) nanobody (e) DARPin (f) F(ab’)_2_ and (g) antibody tLNPs overlaid with NT LNPs. Molar mass (M_w_) profiles of tLNPs did not differ from NT LNPs.

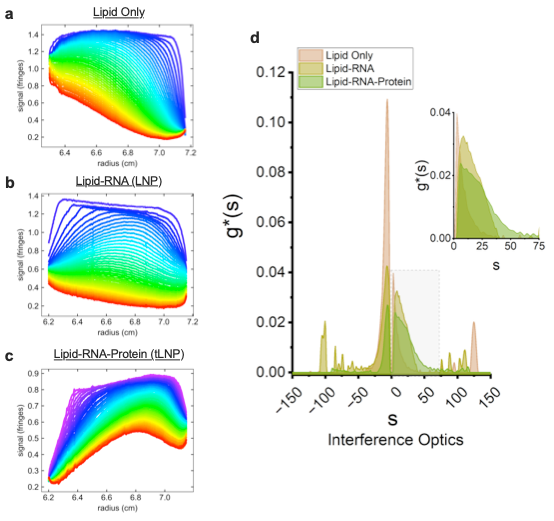

**Supplementary Figure 3 – Sedimentation/Flotation velocity analytical ultracentrifugation (SV/FV-AUC) of antibody-conjugated tLNPs.** Experimental boundaries of (**a**) Lipid only (**b**) Lipid-RNA (NT LNP) and (**c**) Lipid-RNA-Protein (tLNP) by refractive index. (**d**) Lipids alone primarily showed floatation (boundary migration from right to left), while LNPs showed a combination of flotation and sedimentation, likely representing subpopulations differing in RNA content. tLNPs showed increased sedimentation when compared to LNPs, reflecting increases in size and mass after protein functionalization.

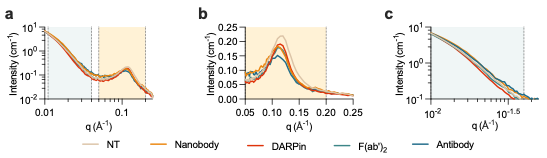

**Supplementary Figure 4 – Comparison of LNP scattering profiles from batch SAXS.** (**a**) Overlaid scattering profiles of NT LNPs and tLNPs depicted between 0.01 < q < 0.2 Å^−1^. The blue region corresponds to the Porod region, while the tan region delineates the Bragg Peak region. (**b**) Zoomed in view from (**a**) showing the Bragg peak region (0.05 < q < 0.20 Å^−1^) of NT LNPs and tLNPs representative of lipid-RNA interactions. tLNPs do not show significant changes in lattice properties compared to NT LNPs, confirming that protein functionalization does not perturb LNP internal structure. (**c**) Zoomed in view from (**a**) showing the Porod region (0.01 < q < 0.04 Å^−1^) of NT LNPs and tLNPs representative of LNP surface area.

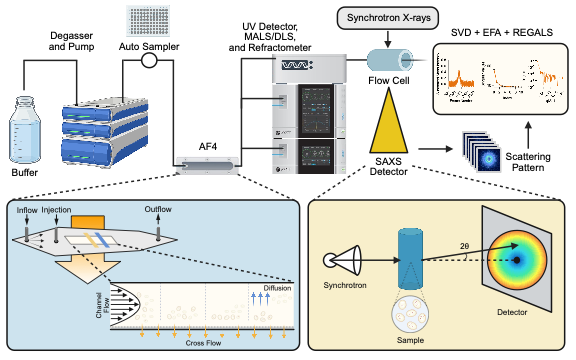
**Supplementary Figure 5 – Overview of AF4-UV-DLS-MALS-SAXS pipeline used to characterize tLNPs.** LNPs were gently separated via AF4 before UV-Vis diode array detector, dynamic light scattering, and multi-angle light scattering analysis. The eluant then entered synchrotron small angle X-ray scattering. To resolve structural species contributing to the SAXS signal across the AF4 elution window, component analysis was performed using SVD and EFA as implemented in REGALS.

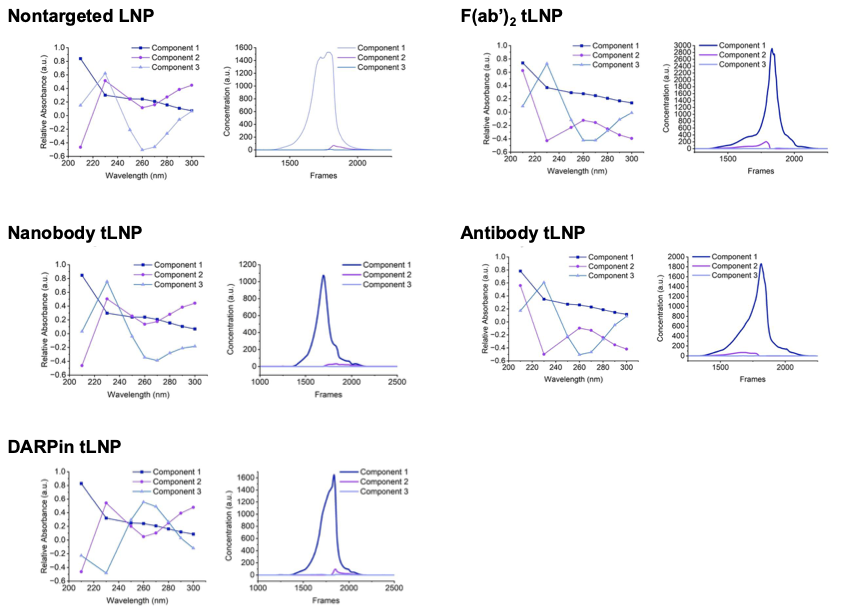

**Supplementary Figure 6 – Multivariate chemometric analysis of the full AF4-UV diode array dataset.** Variation in RNA-associated absorbance across the AF4 elution profile indicates co-existing LNP populations with distinct compositions, motivating chemometric decomposition to resolve overlapping compositional and structural species. Multivariate analysis (SVD, EFA, and MCR-ALS) of AF4–UV diode array data separates lipid-, RNA-, and protein-associated components within LNP and tLNP fractograms, revealing increased compositional heterogeneity and overlapping populations following protein functionalization.

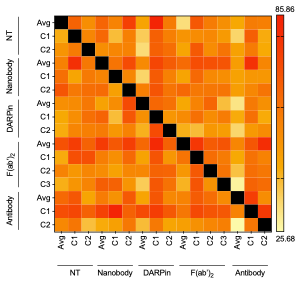

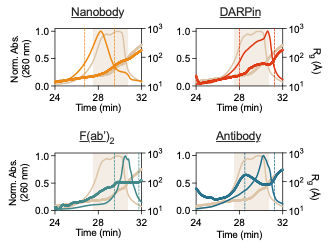

**Supplementary Figure 7 – AF4-MALS derived radius of gyration profiles of LNPs.** R_g_ profiles derived from MALS analysis for NT LNPs (beige) and tLNPs (colors) overlaid with UV fractograms from in-line AF4 separation.

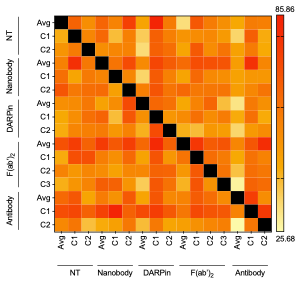

**Supplementary Figure 8 – Volatility of Ratio (V_r_) analysis of SAXS profiles.** Volatility of ratio^2^ (V_r_) analysis quantifies the statistical uniqueness of scattering profiles by measuring deviations between the ensemble-averaged AF4-SAXS signal and SVD-resolved component profiles (C1–C3) across NT LNPs and tLNP formulations. Higher V_r_ values indicate statistically distinct profiles, whereas lower values reflect greater similarity. The observed differences between component-resolved profiles and the ensemble average support the presence of multiple, non-redundant scattering species, consistent with discrete LNP subpopulations rather than a single continuous distribution. These results provide quantitative support for the validity of chemometric decomposition, demonstrating that the resolved components represent statistically independent structural populations rather than mathematical overfitting of the SAXS data.

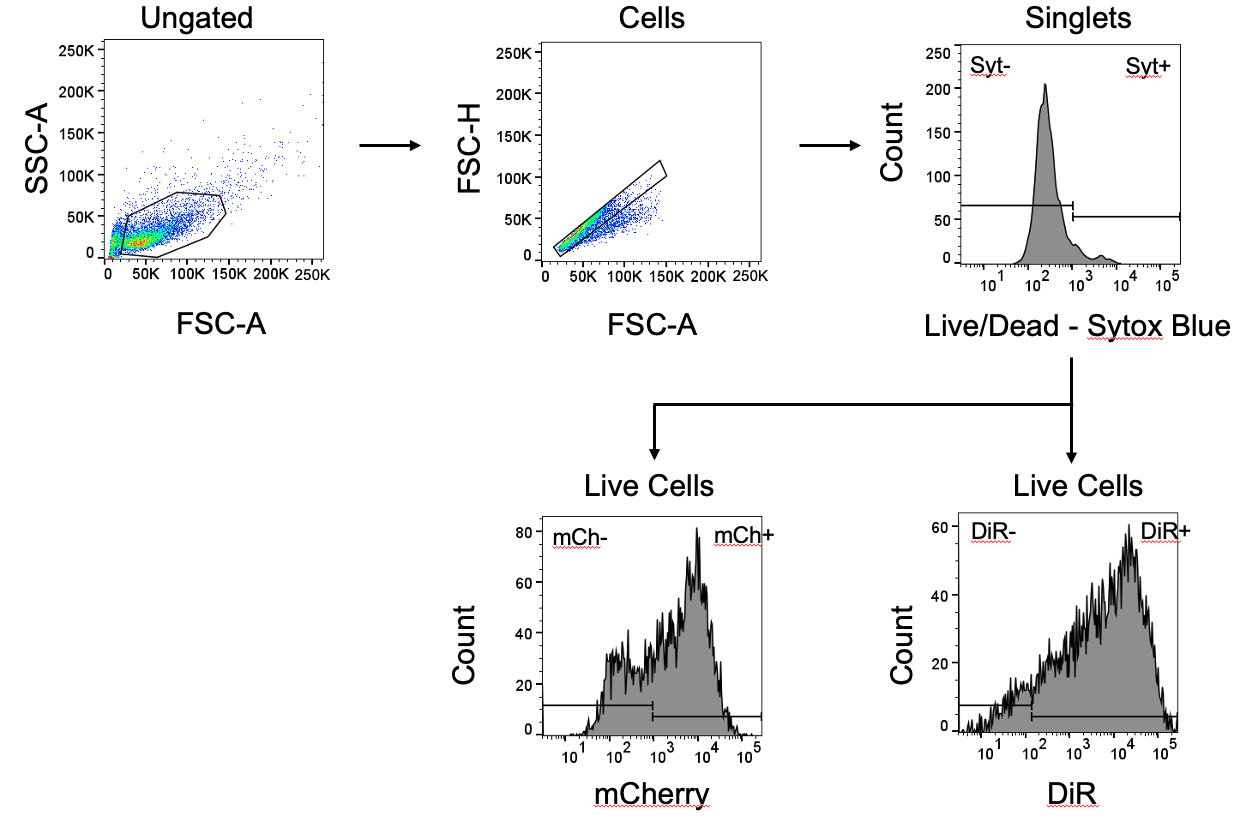

**Supplementary Figure 9 – Representative gating scheme for evaluating LNP accumulation (DiR) and mRNA delivery (mCherry) in BeWo cells.** Data representing DiR and mCherry MFIs are shown in Figure 5 in the main text.

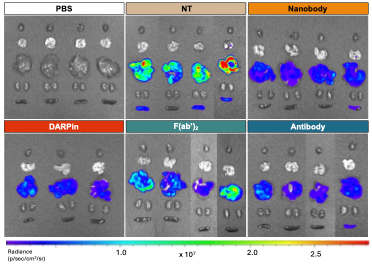

**Supplementary Figure 10 – Luminescence of maternal organs in pregnant mice.** IVIS images of luciferase mRNA delivery (12 µg mRNA per mouse) to the heart, lungs, liver, kidneys, and spleens of pregnant mice from *n = 3-4* biological replicates 6 h following intravenous administration of PBS, NT LNPs, or tLNPs. Images stitched together represent data collected on different days.

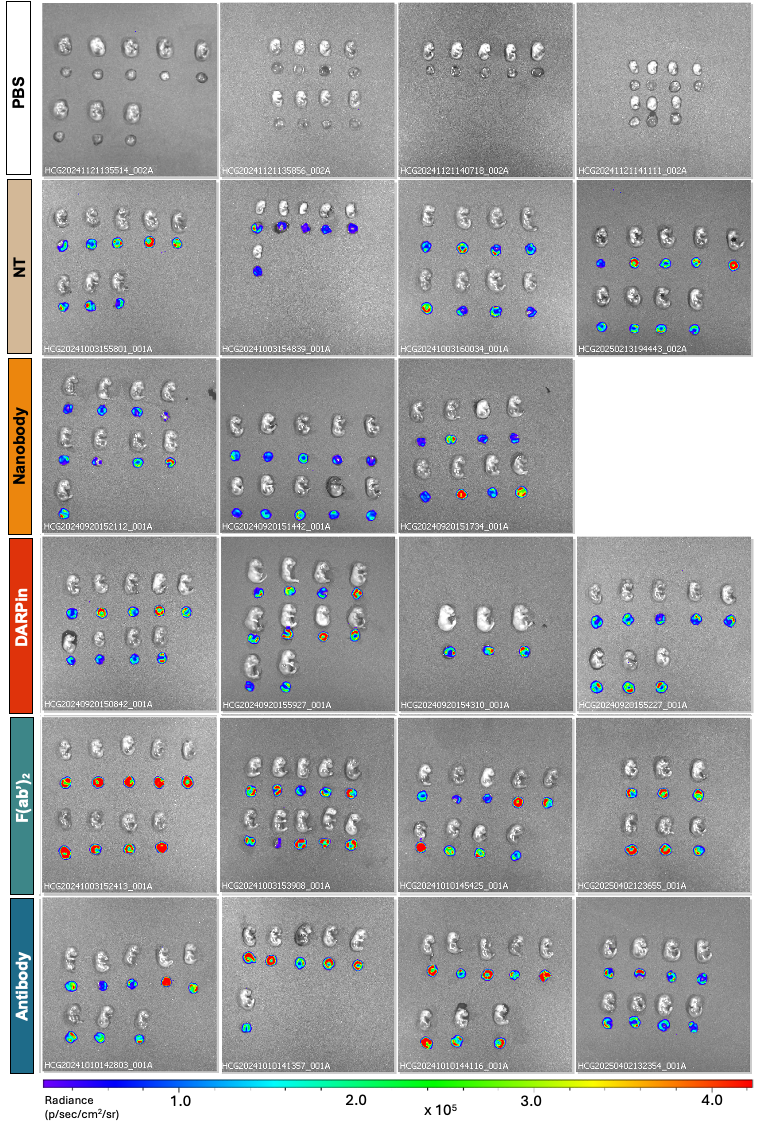

**Supplementary Figure 11 – Luminescence of placentas and fetuses in pregnant mice.** IVIS images of luciferase mRNA delivery (12 µg mRNA per mouse) to the placentas and fetuses of pregnant mice from *n = 3-4* biological replicates 6 h following intravenous administration of PBS, NT LNPs, or tLNPs.

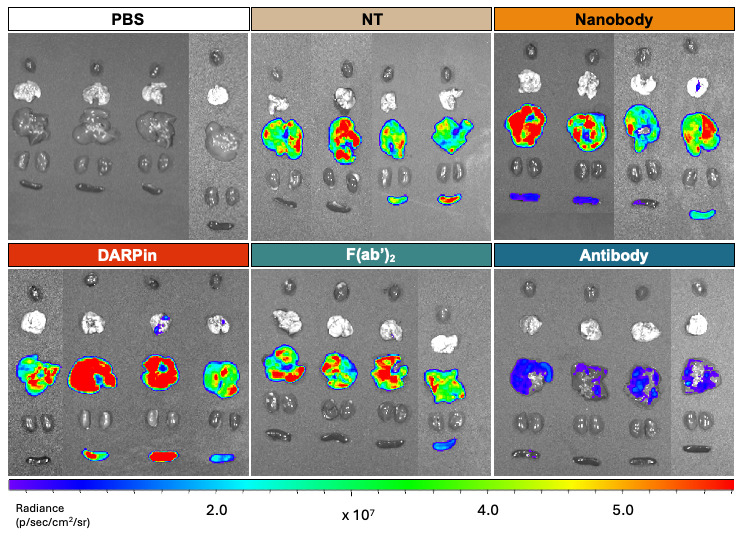

**Supplementary Figure 12 – Luminescence of major organs in nonpregnant mice.** IVIS images of luciferase mRNA delivery (12 µg mRNA per mouse) to the heart, lungs, liver, kidneys, and spleens of nonpregnant mice from *n = 4* biological replicates 6 h following intravenous administration of PBS, NT LNPs, or tLNPs. Images stitched together represent data collected on different days.

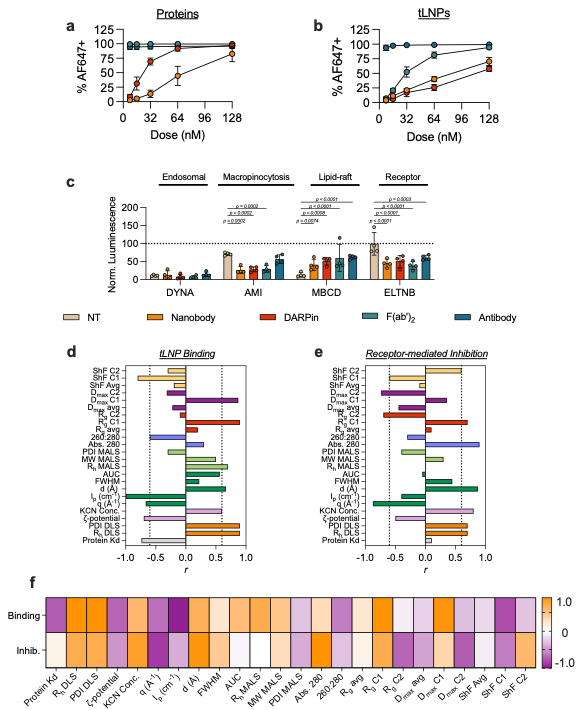

**Supplementary Figure 13 – tLNP binding, endocytosis, and subsequent correlations to biophysical parameters.** Alexafluor647 (AF647)-labeled Nanobody, DARPin, F(ab’)_2_, and antibody (**a**) proteins and (**b**) tLNPs were incubated with BeWo placental cells at varying doses for 1 h before analysis via flow cytometry. (**c**) Various pathways of endocytosis were inhibited using small molecule inhibitors (DYNA – dynasore; AMI – amiloride; MBCD – methyl-beta-cyclodextran; ELTNB – erlotinib) in BeWo placental cells for 1 h before dosing with NT LNPs or tLNPs. After 24 h, luminescence was measured and values were normalized to cellular viability. NT LNPs were primarily inhibited via MBCD, while tLNPs were inhibited using AMI and ELTNB. tLNP binding from (**b**) and ELNTB inhibition from (**c**) were then correlated with biophysical parameters using Spearman Correlation (**d-f**). Both binding and inhibition had positive correlation with DLS parameters and KCN protein quantification. tLNP binding had a negative correlation with protein Kd values and the 260:280 ratio. Binding and inhibition had inverse correlation patterns regarding R_g_, D_max_, and shape factor. Binding correlated positively with R_g_ and D_max_ of C1 and negatively with shape factor C1, while inhibition showed inverse correlations.

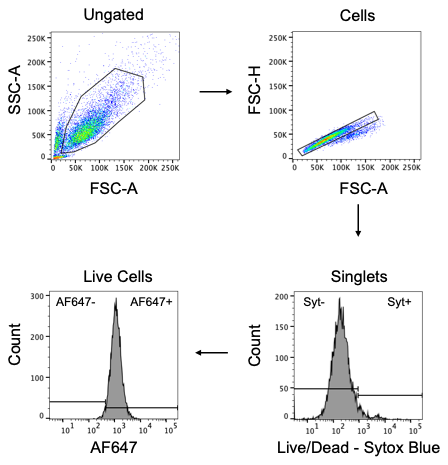

**Supplementary Figure 14 – Representative gating strategy for evaluating AF647-labeled protein and tLNP binding in BeWo cells.** Data representing % AF647+ populations are shown in Supplementary Figure 11.

**SUPPLEMENTARY TABLES**

**Supplementary Table 1 – Characterization of LNPs using traditional characterization techniques.**

| **LNP** | **R_h_ (nm)** | **PDI** | **Zeta potential (mV)** | **Protein**  **conc.**  **(ng µL^-1^)** | **Encapsulation efficiency (%)** | **mRNA conc.**  **(ng µL^-1^)** | **Particle conc. per mL** |
| --- | --- | --- | --- | --- | --- | --- | --- |
| NT | 45.4 ± 25.8 | 0.32 ± 0.04 | 9.86 ± 0.4 | 0.00 ± 3.7 | 99.3 ± 0.8 | 40.8 ± 6.3 | 2.05E12 |
| Nanobody | 51.4 ± 29.2 | 0.28 ± 0.03 | -4.79 ± 0.6 | 10.32 ± 8.5 | 99.6 ± 0.3 | 35.2 ± 0.9 | 9.29E11 |
| DARPin | 50.7 ± 28.8 | 0.26 ± 0.04 | 2.63 ± 1.4 | 3.00 ± 2.4 | 97.0 ± 0.1 | 30.3 ± 4.5 | 5.72E11 |
| F(ab')_2_ | 69.0 ± 39.2 | 0.30 ± 0.06 | 1.31 ± 0.1 | 8.54 ± 1.4 | 97.2 ± 0.5 | 44.9 ± 1.8 | 1.86E12 |
| Antibody | 80.8 ± 50.0 | 0.25 ± 0.02 | -0.20 ± 1.4 | 8.29 ± 8.6 | 97.3 ± 0.6 | 35.7 ± 4.5 | 3.51E12 |

**Supplementary Table 2 – Calculated properties of tLNPs for calculation of dn/dc.**

| **Construct** | **Added protein mass (MDa)** | **New total mass (MDa)** | **Lipid mass fraction** | **RNA mass fraction** | **Protein mass fraction** | **dn/dc (mL/g)** |
| --- | --- | --- | --- | --- | --- | --- |
| NT LNP | 0 | 57.94 | 0.9828 | 0.0172 | 0 | 0.16 |
| +15.8 kDa Nanobody (×90) | 1.422 | 59.362 | 0.9599 | 0.0168 | 0.0241 | 0.161 |
| +17.9 kDa DARPin (×90) | 1.611 | 59.551 | 0.9727 | 0.0168 | 0.0271 | 0.161 |
| +110 kDa  F(ab’)_2_ (×90) | 9.900 | 67.84 | 0.8391 | 0.0147 | 0.1459 | 0.165 |
| +150 kDa Antibody (×90) | 13.500 | 71.44 | 0.7979 | 0.014 | 0.1889 | 0.165 |

**Supplementary Table 3 – Table of parameters derived from Multiple Lorentz fitting of batch SAXS data.**

|  | Fit Deconvoluted H^\|\|^ Peak | | | | | Fit Disorder Region | | | | |  |
| --- | --- | --- | --- | --- | --- | --- | --- | --- | --- | --- | --- |
| Sample | q (Å^-1^) | I_p_ (cm^-1^) | d (Å) | Area | FWHM | q (Å^-1^) | I_p_ (cm^-1^) | d (Å) | Area | FWHM | Area^H\|\|^/  Area^Disorder^ |
| NT | 0.116 | 0.377 | 54.1 | 0.028 ± 0.001 | 0.046 ± 0.001 | 0.062 | 0.075 | 101.3 | 0.023 ± 0.008 | 0.195 ± 0.040 | 1.2 |
| Nanobody | 0.111 | 0.390 | 56.6 | 0.016 ± 0.001 | 0.044 ± 0.002 | 0.060 | 0.115 | 104.7 | 0.024 ± 0.006 | 0.143 ± 0.022 | 0.7 |
| DARPin | 0.113 | 0.385 | 55.6 | 0.026 ± 0.001 | 0.044 ± 0.001 | 0.061 | 0.067 | 103.0 | 0.004 ± 0.001 | 0.052 ± 0.010 | 6.5 |
| F(ab')_2_ | 0.113 | 0.223 | 55.6 | 0.014 ± 0.001 | 0.042 ± 0.002 | 0.060 | 0.061 | 104.7 | 0.006 ± 0.003 | 0.084 ± 0.036 | 2.3 |
| Antibody | 0.112 | 0.100 | 56.1 | 0.006 ± 0.001 | 0.041 ± 0.003 | 0.063 | 0.074 | 99.7 | 0.019 ± 0.004 | 0.161 ± 0.024 | 0.3 |

**Supplementary Table 4 – Table of Parameters derived from in-line DLS-MALS analysis.**

|  |  | NT | Nanobody | DARPin | F(ab')_2_ | Antibody |
| --- | --- | --- | --- | --- | --- | --- |
| In-line DLS | **R_h_ (nm)** | 28.1 ± 0.5% | 28.1 ± 0.4% | 30.1 ± 0.6% | 38.8 ± 2.2% | 50.2 ± 1.0% |
| MALS | **M_n_ (MDa)** | 39.6 ± 3.1% | 35.8 ± 0.6% | 68.3 ± 7.9% | 110 ± 12.1% | 162 ± 23.4% |
|  | **M_w_ (MDa)** | 46.5 ± 6.2% | 39.8 ± 0.9% | 79.7 ± 13.1% | 112 ± 12.0% | 168 ± 21.1% |
|  | **Mw/Mn** | 1.2 ± 6.9% | 1.1 ± 1.0% | 1.2 ± 15.3% | 1.0 ± 17.1% | 1.0 ± 31.5% |
|  | **R_g_ (Å)** | 400 ± 13.1% | 269 ± 3.7% | 505 ± 15.0% | 942 ± 5.0% | 960 ± 8.0% |

**Supplementary Table 5 – Porod Exponents derived from batch SAXS and AF4-UV-DLS-MALS-SAXS profiles.**

|  | NSLSII LiXS | APS Biocat | | | |
| --- | --- | --- | --- | --- | --- |
|  | Batch | Avg | C1 | C2 | C3 |
| NT | 3.6 | 3.6 | 3.8 | 3.2 |  |
| Nanobody | 3.6 | 3.0 | 3.2 | 3.9 |  |
| DARPin | 4.0 | 3.3 | 3.4 | 3.1 |  |
| F(ab’)_2_ | 3.6 | 3.7 | 3.1 | 3.6 | 3.8 |
| Antibody | 3.3 | 3.1 | 2.5 | 3.9 |  |

**Supplementary Table 6 – Table of determined parameters from AF4-SAXS with SVD-EFA analyses.**

| Sample |  | Guinier | | GNOM | |
| --- | --- | --- | --- | --- | --- |
|  |  | qR_g_ | R_g_ (Å) | R_g_ (Å) | D_max_ (Å) (Å) |
| NT | Avg | 0.74 – 1.55 | 266.6 ± 9.3 | 269.8 ± 5.4 | 824 |
|  | C1 | 0.49 – 1.29 | 132.4 ± 18.4 | 162.2 ± 52 | 661 |
|  | C2 | 0.71 – 1.83 | 284.6 ± 8.6 | 325.1 ± 9.4 | 1040 |
| Nanobody | Avg | 0.62 – 1.45 | 225.0 ± 14.5 | 218.8 ± 17.1 | 716 |
|  | C1 | 0.61 – 1.68 | 188.6 ± 8.0 | 210.7 ± 15.2 | 714 |
|  | C2 | 0.66 – 1.54 | 219.9 ± 13.4 | 222.8 ± 7.5 | 716 |
| DARPin | Avg | 0.49 – 1.36 | 178.1 ± 5.3 | 182.3 ± 5.3 | 629 |
|  | C1 | 0.53 – 1.30 | 175.9 ± 11.6 | 186.7 ± 3.8 | 515 |
|  | C2 | 0.46 – 1.40 | 177.3 ± 7.7 | 189.8 ± 9.3 | 714 |
| F(ab’)_2_ | Avg | N/A | N/A | 533.9 ± 8.91 | 1507^A^ |
|  | C1 | N/A | N/A | 303.5 ± 1.66 | 717 |
|  | C2 | N/A | N/A | N/A | 1677 ^A^ |
|  | C3 | 0.55 – 1.32 | 190.4 ± 18.1 | 202.5 ± 22.2 | 714 |
| Antibody | Avg | N/A | N/A | 226.4 ± 11.4 | 716 |
|  | C1 | N/A | N/A | 295.5 ± 2.1 | 717 |
|  | C2 | 0.70 – 1.30 | 232.9 ± 18.4 | 239.8 ± 3.8 | 716 |

A. For these largest species, the Shannon criterion (q_min_·D_max_ ≳ π) was not fully satisfied, indicating limited information content at the longest real-space distances. In these cases, interpretation was restricted to short- and intermediate-range features of the P(r) and to comparative parameters such as R_g_, while absolute D_max_ values obtained are viewed as lower bounds.

**Supplementary Table 7 – Shape Factor (D_max_ / R_g_) from P(r) analyses.**

| **Sample** | **Component** | **Shape Factor** |
| --- | --- | --- |
| NT | Avg | 3.05 |
|  | C1 | 4.07 |
|  | C2 | 3.20 |
| Nanobody | Avg | 3.27 |
|  | C1 | 3.38 |
|  | C2 | 3.21 |
| DARPin | Avg | 3.45 |
|  | C1 | 2.75 |
|  | C2 | 3.76 |
| F(ab’)_2_ | Avg | 2.82 |
|  | C1 | 2.36 |
|  | C2 | N/A |
|  | C3 | 3.52 |
| Antibody | Avg | 3.16 |
|  | C1 | 2.42 |
|  | C2 | 2.98 |

**Supplementary Table 8 – Determined size parameters from DENSS ab initio reconstructions.**

| Sample | Component ^A^ | Length (X) (Å) | Width (Y) (Å) | Height (Z) (Å) |
| --- | --- | --- | --- | --- |
| NT | Avg | 587.8 | 411.4 | 352.7 |
|  | C2 | 585.0 | 780.0 | 828.8 |
|  | C1 | 309.8 | 216.9 | 340.8 |
| Nanobody | Avg | 605.3 | 330.2 | 440.3 |
|  | C2 | 469.9 | 369.2 | 704.8 |
|  | C1 | 435.1 | 669.4 | 368.2 |
| DARPin | Avg | 614.6 | 279.4 | 279.4 |
|  | C1 | 434.5 | 507.0 | 458.7 |
|  | C2 | 301.2 | 368.2 | 635.9 |
| F(ab’)_2_ | C1 | 1478.5 | 1196.9 | 1408.1 |
|  | C3 | 535.5 | 535.5 | 301.2 |
|  | Avg | 474.8 | 593.4 | 415.4 |
|  | C2 | N/A | N/A | N/A |
| Antibody | C1 | 739.4 | 739.4 | 571.4 |
|  | C2 | 503.4 | 704.8 | 436.3 |
|  | Avg | 426.0 | 266.3 | 479.3 |

A. Components are sorted longest to shortest by length (X) within each LNP formulation.

**Supplementary Table 9 – Two-tailed P values from Spearman Correlations to luminescence.**

|  | **Placental** | **Preg. Hepatic** | **Nonpreg. Hepatic** |
| --- | --- | --- | --- |
| Protein Kd | -0.9487 | 0.1054 | 0.9487 |
| R_h_ DLS | 0.6 | -0.6 | -0.6 |
| PDI DLS | -0.2 | 1 | 0.2 |
| ζ-potential | 0.1 | 0.5 | -0.1 |
| KCN Conc. | 0 | -0.2 | 0.2 |
| q (Å^-1^) | -0.2052 | 0.1539 | -0.0513 |
| I_p_ (cm^-1^) | -0.6 | 0.6 | 0.6 |
| d (Å) | 0.2052 | -0.1539 | 0.0513 |
| FWHM | 0.2236 | -0.6708 | 0.2236 |
| AUC | 0.8721 | -0.0513 | -0.9747 |
| R_h_ In-line DLS | 0.8721 | -0.6156 | -0.8208 |
| R_g_ MALS | 0.9 | -0.5 | -0.9 |
| MW MALS | 0.9 | -0.5 | -0.9 |
| PDI MALS | -0.6325 | 0.3162 | 0.6325 |
| Abs. 280 | -0.1 | -0.1 | 0.5 |
| 260:280 | -1 | 0.2 | 0.8 |
| R_g_ avg | 0.5 | 0.6 | -0.6 |
| R_g_ C1 | 0.7 | -0.3 | -0.5 |
| R_g_ C2 | -0.3 | 0.3 | -0.3 |
| D_max_ avg | -0.2236 | 0.6708 | -0.2236 |
| D_max_ C1 | 0.5643 | -0.1539 | -0.6669 |
| D_max_ C2 | -0.5798 | 0.3162 | 0 |
| ShF Avg | -0.5 | -0.6 | 0.6 |
| ShF C1 | -0.9 | 0.4 | 0.6 |
| ShF C2 | 0.1 | 0.1 | 0.5 |

**Supplementary Table 10 – Two-tailed P values from Spearman Correlations to toxicity in pregnant mice.**

|  | **Binding** | **Inhibition** |
| --- | --- | --- |
| Protein Kd | -0.7379 | 0.1054 |
| R_h_ DLS | 1 | 0.4 |
| PDI DLS | -0.6 | 0 |
| ζ-potential | -0.7 | -0.5 |
| KCN Conc. | 0.6 | 0.8 |
| q (Å^-1^) | -0.6669 | -0.8721 |
| I_p_ (cm^-1^) | -1 | -0.4 |
| d (Å) | 0.6669 | 0.8721 |
| FWHM | 0.2236 | 0.4472 |
| AUC | 0.5643 | -0.0513 |
| R_h_ In-line DLS | 0.8208 | 0.1539 |
| R_g_ MALS | 0.7 | 0 |
| MW MALS | 0.7 | 0 |
| PDI MALS | -0.9487 | -0.4743 |
| Abs. 280 | 0.3 | 0.9 |
| 260:280 | -0.6 | -0.3 |
| R_g_ avg | 0.2 | 0.1 |
| R_g_ C1 | 0.9 | 0.7 |
| R_g_ C2 | -0.1 | -0.7 |
| D_max_ avg | -0.2236 | -0.4472 |
| D_max_ C1 | 0.8721 | 0.3591 |
| D_max_ C2 | -0.3162 | -0.7379 |
| ShF Avg | -0.2 | -0.1 |
| ShF C1 | -0.8 | -0.6 |
| ShF C2 | -0.3 | 0.6 |

**Supplementary Table 11 – Two-tailed P values from Spearman Correlations to toxicity in pregnant mice.**

|  | **C3a** | **TNF** | **IL6** | **IFNg** | **ALT** | **AST** |
| --- | --- | --- | --- | --- | --- | --- |
| Protein Kd | 0.3162 | -0.9487 | 0.2108 | -0.2108 | 0.6325 | 0.2108 |
| R_h_ DLS | 0.3 | 0.9 | 0.3 | 0.8 | -0.2 | -0.6 |
| PDI DLS | -0.1 | -0.7 | -0.9 | -0.4 | 0.6 | -0.2 |
| ζ-potential | -0.7 | -0.4 | -0.3 | -0.8 | -0.3 | 0.6 |
| KCN Conc. | 0.5 | 0.3 | 0.1 | 0.6 | 0.6 | -0.8 |
| q (Å^-1^) | -0.3591 | -0.4104 | -0.0513 | -0.5643 | -0.5643 | 0.8721 |
| I_p_ (cm^-1^) | -0.3 | -0.9 | -0.3 | -0.8 | 0.2 | 0.6 |
| d (Å) | 0.3591 | 0.4104 | 0.0513 | 0.5643 | 0.5643 | -0.8721 |
| FWHM | -0.4472 | 0.4472 | 0.8944 | -0.2236 | -0.2236 | 0.2236 |
| AUC | -0.2052 | 0.6669 | -0.2052 | 0.3078 | -0.4617 | -0.3078 |
| R_h_ In-line DLS | -0.2052 | 0.9747 | 0.4104 | 0.4104 | -0.6156 | -0.2052 |
| R_g_ MALS | -0.3 | 0.9 | 0.3 | 0.3 | -0.7 | -0.1 |
| MW MALS | -0.3 | 0.9 | 0.3 | 0.3 | -0.7 | -0.1 |
| PDI MALS | -0.3162 | -0.7906 | 0 | -0.7906 | 0 | 0.7906 |
| Abs. 280 | 0.2 | 0.1 | 0.2 | 0.2 | 0.7 | -0.6 |
| 260:280 | 0.5 | -0.8 | -0.1 | -0.1 | 0.4 | 0.3 |
| R_g_ avg | 0 | 0.1 | -0.8 | 0.2 | 0.2 | -0.6 |
| R_g_ C1 | 0.1 | 0.8 | 0.1 | 0.6 | 0.1 | -0.8 |
| R_g_ C2 | 0.6 | -0.3 | -0.6 | 0.4 | -0.1 | 0 |
| D_max_ avg | 0.4472 | -0.4472 | -0.8944 | 0.2236 | 0.2236 | -0.2236 |
| D_max_ C1 | 0.4104 | 0.6669 | -0.2052 | 0.8208 | 0.0513 | -0.8208 |
| D_max_ C2 | 0.0513 | -0.0513 | -0.8721 | 0.1539 | 0.4104 | -0.6669 |
| ShF Avg | 0 | -0.1 | 0.8 | -0.2 | -0.2 | 0.6 |
| ShF C1 | 0.3 | -0.9 | -0.3 | -0.3 | 0.2 | 0.5 |
| ShF C2 | -0.7 | -0.1 | 0.3 | -0.7 | 0.3 | 0.1 |

**Supplementary Table 12 – Two-tailed P values from Spearman Correlations to toxicity in nonpregnant mice.**

|  | **C3a** | **TNF** | **IL6** | **IFNg** | **ALT** | **AST** |
| --- | --- | --- | --- | --- | --- | --- |
| Protein Kd | 0.9487 | -0.7379 | -0.9487 | -0.6325 | -0.7379 | -0.6325 |
| R_h_ DLS | -0.3 | 1 | 0.7 | 0.7 | 1 | 0.1 |
| PDI DLS | 0.1 | -0.6 | -0.5 | -0.9 | -0.6 | -0.3 |
| ζ-potential | -0.3 | -0.7 | 0 | -0.3 | -0.7 | 0.4 |
| KCN Conc. | 0.5 | 0.6 | -0.1 | 0.1 | 0.6 | -0.7 |
| q (Å^-1^) | -0.4104 | -0.6669 | -0.0513 | -0.1539 | -0.6669 | 0.6669 |
| I_p_ (cm^-1^) | 0.3 | -1 | -0.7 | -0.7 | -1 | -0.1 |
| d (Å) | 0.4104 | 0.6669 | 0.0513 | 0.1539 | 0.6669 | -0.6669 |
| FWHM | 0.4472 | 0.2236 | 0.2236 | 0.6708 | 0.2236 | -0.2236 |
| AUC | -0.8208 | 0.5643 | 0.8721 | 0.4104 | 0.5643 | 0.5643 |
| R_h_ MALS | -0.5643 | 0.8208 | 0.9747 | 0.8721 | 0.8208 | 0.4617 |
| MW MALS | -0.7 | 0.7 | 1 | 0.8 | 0.7 | 0.6 |
| PDI MALS | -0.7 | 0.7 | 1 | 0.8 | 0.7 | 0.6 |
| Abs. 280 | 0.3162 | -0.9487 | -0.6325 | -0.4743 | -0.9487 | 0 |
| 260:280 | 0.8 | 0.3 | -0.3 | 0 | 0.3 | -0.9 |
| R_g_ avg | 0.5 | -0.6 | -0.9 | -0.6 | -0.6 | -0.3 |
| R_g_ C1 | -0.5 | 0.2 | 0.3 | -0.3 | 0.2 | 0.1 |
| R_g_ C2 | -0.1 | 0.9 | 0.6 | 0.5 | 0.9 | -0.2 |
| D_max_ avg | -0.6 | -0.1 | -0.1 | -0.4 | -0.1 | 0.5 |
| D_max_ C1 | -0.4472 | -0.2236 | -0.2236 | -0.6708 | -0.2236 | 0.2236 |
| D_max_ C2 | -0.4104 | 0.8721 | 0.5643 | 0.3078 | 0.8721 | 0.0513 |
| ShF Avg | -0.3078 | 0.1026 | 0.1026 | -0.4617 | 0.1026 | -0.1026 |
| ShF C1 | 0.5 | -0.2 | -0.3 | 0.3 | -0.2 | -0.1 |
| ShF C2 | 0.2 | -0.8 | -0.8 | -0.7 | -0.8 | 0 |
